# Unravelling the Genetic Architecture of Field Traits through Multi-Omics Platform Data integration

**DOI:** 10.1101/2025.10.27.684747

**Authors:** Baber Ali, Stéphane Nicolas, Mélisande Blein-Nicolas, Marie-Laure Martin, Yacine Djabali, Tristan Mary-Huard, Alain Charcosset, Laurence Moreau, Renaud Rincent

## Abstract

**Background:** Identifying the genes and regulatory regions underlying a complex trait is a long-standing challenge. GWAS is generally used, but it suffers from insufficient power, lack of resolution and inefficiency to systematically screen for epistasis. We propose a systems genetics approach integrating multi-omics to overcome these limits. It was applied to a panel of maize with the objective to analyze the genetic determinism of yield in a multi-environment trials by combining genomics with transcriptomics measured on a platform.

**Results:** Despite the contrasted conditions between the platform and the fields, transcriptomics could be used to identify candidate genes. A presence-absence variation was in particular detected, and the transcripts allowed the identification of causal genes, increasing resolution in comparison to GWAS. In total, 47 genes were identified along the genome, and we could characterize their contrasted effect on yield according to environmental covariates. We demonstrated that the cis- and also the trans-eQTLs of these genes had an important contribution to genetic variance, suggesting a key role of epistatic interactions. In terms of predictive ability, the cis-eQTL resulted in an increase of 39 to 52% on average across the environments, in comparison to random SNP sets.

**Conclusions:** By efficiently combining multi-omics, it is possible to considerably increase our understanding of genetic architecture in comparison to standard GWAS. We demonstrated that omics data even measured on a phenotyping platform can be used for the analysis of field traits, opening the way for their routine use in plant breeding both for marker-assisted selection and bio-informed predictions.

## Background

Agriculture production systems are facing higher temperatures and changes in rain patterns because of climate change. Most of the agriculture is rainfed, and changes in weather patterns are challenging the usual schedules of sowing, maturation, and harvesting of several crops, including maize. A study using several crop models has projected a decrease of up to 24% in global maize production by the end of this century (1). Therefore, constant genetic improvement of maize, the world’s most cultivated plant species, is necessary to meet the sustainable development goal of global food security set by the United Nations (UN) (2). Understanding the genetic architecture of the target traits is necessary so that the underlying key genomic regions can be capitalized on in breeding programs using marker-assisted selection or bio-informed genomic predictions.

Genome-wide association study (GWAS) is one such approach that helps identify key quantitative trait loci (QTLs) associated with a trait (3). In GWAS, QTLs are detected based on the linkage disequilibrium (LD) between the causal variant and surrounding variants. As we cannot characterize all polymorphisms, LD is essential to capture the effect of any causal polymorphism. The downside is that LD can be quite strong within a given region, making it difficult to identify the causal variant. Another important limit of GWAS is that the very large number of polymorphisms make it impossible to systematically screen for epistatic relationships, in particular because of the multiple testing burden. This is a severe drawback, as we know that genes interact in complex networks, with a fundamental role of regulatory elements (4). Under such circumstances, we need complementary approaches, more resolutive and more powerful, and that would be able to detect such regulatory and epistatic interactions.

Transcriptome-wide association studies (TWAS) test the associations between transcript abundances and the target phenotype. As a consequence, it directly identifies underlying genes, leading to a very high resolution (5). The downside being that regions outside genes are not evaluated. By combining phenotypic data collected from a single location and transcriptomic data collected on the same plants, studies have shown that TWAS can complement GWAS in dissecting the architecture of traits in several plant species, such as maize (6,7), cotton (8), and soybean (9). In most omics-based studies, plants are grown on a platform to better control the environment and ease the sampling process. In these studies, omics are then used to investigate the determinism of traits measured under the same conditions, and most often on the same plants. But, even if plants grown on platforms face different environmental conditions than those in the fields, different studies have proven that platform omics could be predictive of traits measured in the fields (10,11). Demonstrating the interest of platform omics to decipher field traits would be of considerable interest to the genetic and plant breeding communities, as the costs for omics characterization still prevent us from using it systematically in each field trial. Our study offers novelty as it uses transcriptomic data measured in controlled conditions of a platform to model productivity in a multi-environment trial (MET), which according to our knowledge has not been explored so far. In this context, we evaluated if platform omics are useful for dissecting the genetic architecture of complex field traits in the presence of genotype x environment (GxE) interactions.

Maize grain yield is generally measured in METs across different years and locations to evaluate the genotype performances in contrasted conditions. In these METs, each genotype may have a particular response to the different environmental conditions, leading to GxE interactions. This means that the QTL or gene response differs from one environment to another (12). For an efficient use of these regions in breeding, this clearly needs to be considered in association studies. One first approach is to estimate reaction norms of genotypes against environmental covariates, such as temperature, soil water potential or radiation intensity. Then, associations between genome-wide markers and the reaction norms are evaluated to identify key genomic regions (13). Another approach is to independently perform association analysis in each environment (12,14,15), or set of similar environments (16). However, interpreting the results of multiple association studies can become quite challenging, especially when the number of environments is high. Therefore, a meta-analysis of the results of association studies can be a practical approach to summarize individual environment results and increase detection power and interpretability. Recently, new methods have been proposed to perform meta-analyses in MET (17). The authors of this method introduced two meta-analysis procedures: fixed effect (FE) and random effect (RE). FE procedure considers that a marker effect is constant for all the environments or at least a group of similar environments. RE considers these effects to be random, i.e., either equal or different from one environment to another. Both procedures have been found to be effective for the meta-analysis of GWAS, but they have never been applied to summarize the results of TWAS in a multi-environmental setting.

In the present study, we used a transcriptomic dataset obtained under controlled conditions from a maize panel of 244 genotypes to identify its ability to explain the genetic architecture of a complex quantitative trait, grain yield, measured in field conditions in 25 trials. By combining TWAS and eQTL analysis, we demonstrate that SNPs and transcriptomics data can be used complementarily to identify causal genes as well as numerous regulatory regions potentially responsible of epistatic relationships, that were undetected using GWAS. Importantly, these identified cis and trans regions considerably contribute to genetic variance and improve genomic prediction accuracy in comparison to SNPs randomly sampled along the genome. The integration of these results using meta-analysis allowed us to explain the gene x environment interactions using environmental covariates.

## Results

### Genome-Wide Associations Reveal Key Genomic Regions Associated with Yield

GWAS was performed to identify genomic regions associated with grain yield in a panel of 244 maize hybrids grown in 25 field trials (15). GWAS was performed on each of the 25 trials using a unified mixed model (18). It resulted in one GWAS per field trial, and all the resulting GWAS were then subjected to meta-analysis (17) to increase detection power and interpretability of the results.

Based on a false discovery rate (FDR) of 10%, two peaks were identified with the fixed effect (FE) procedure on the GWAS results (Fig. 1A): on chromosome 3 (127 Mb to 158 Mb) and on chromosome 6 (14 Mb to 23 Mb). Based on SNP coordinates on the AGPv4 maize genome assembly, using a 10 Kb window upstream and downstream of all these significant SNPs, we identified 15 unique genes (Table S1). Similarly, for the random effect (RE) procedure (Fig. 1B), we identified nine significant peaks on chromosomes 1 (285 Mb), 2 (134 Mb), 3 (48 Mb), 8 (13 Mb), and 10 (133.4 Mb to 133.7 Mb) and three peaks on chromosome 6 (7 Mb to 7.5 Mb, 14 Mb to 16 Mb, 21.2 Mb to 25.2 Mb). Using a 10 Kb window upstream and downstream of these SNPs, 41 unique genes were identified (Table S2). Combining both gene sets resulted in 44 unique genes. Gene ontology (GO) analysis using agriGO V2.0 (19) identified an enrichment in genes (5 out of 44) related to ubiquitin-protein transferase activity molecular function (Fig S1), which is known to be associated with abiotic stress response in maize (20). This molecular function regulates several biological processes, signal transduction, and protein interactions.

**Fig. 1.**
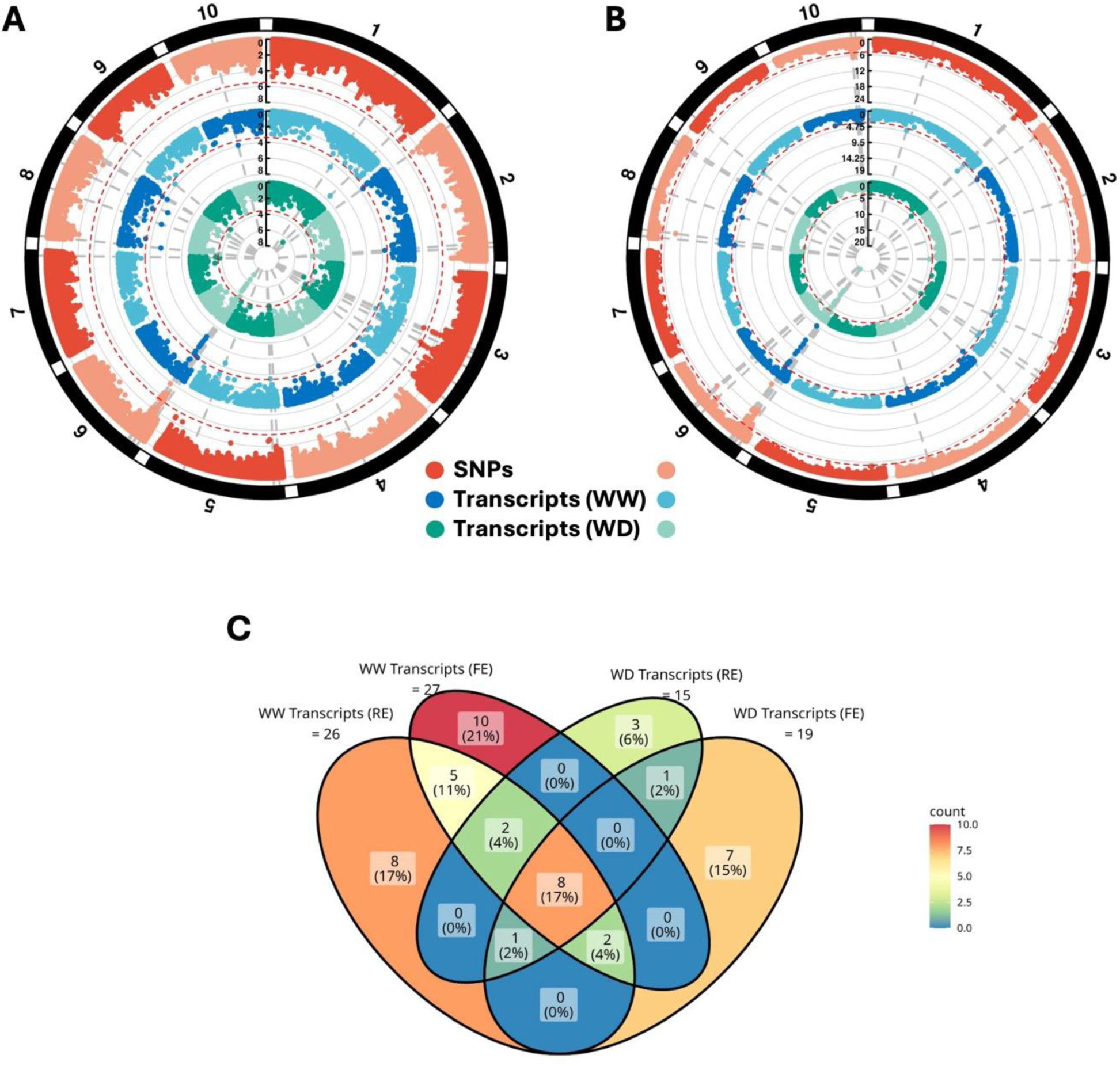
GWAS and TWAS on grain yield. A) Fixed Effect (FE) procedure of meta-analysis. B) Random Effect (RE) procedure of meta-analysis. Numbers 1 to 10 represent different chromosomes. Red dots represent the GWAS (SNPs) Manhattan plot, blue dots represent TWAS-WW, and green dots represent TWAS-WD. Red dashed lines represent the FDR (10%) significance threshold. Grey dashed lines represent the presence of significant signals. The scale bar from 0 to 8 represents the −log_10_ of p-values. C) Venn diagram of the numbers of significantly associated genes obtained with TWAS with the fixed effect (FE) and random effect (RE) meta-analysis procedures. The color scale represents the number of genes within each group.

### Transcriptome-Wide Associations Directly Identify Genes Associated with Yield

Transcriptomic data was acquired in 2013 on the same 244 genotypes in controlled conditions (phenotyping platform) in two contrasting watering regimes, i.e., water deficit (WD) and well-watered (WW). Similarly to the approach applied for GWAS, we tested the association between the transcripts measured in both water conditions and grain yield in each field (TWAS-WW and TWAS-WD), followed by a meta-analysis.

In the case of TWAS-WW, 27 and 26 genes were found to be significantly associated (10% FDR nominal value) with yield using FE and RE procedures, respectively (Fig. 1). Similarly, for TWAS-WD, 19 and 15 genes were significantly associated (10% FDR nominal value) with yield using FE and RE procedures, respectively. All these four sets of genes correspond to 47 unique genes (Table S3). Eight genes were found to be common among these four sets, suggesting their effects to be stable across both watering conditions (Fig. 1C). These common genes are *Zm00001d034045, Zm00001d035346, Zm00001d035367, Zm00001d035377, Zm00001d035390, Zm00001d035400, Zm00001d035409, Zm00001d035420*. Five of these genes were described in the AGPv4 assembly of the maize genome (Table 1), while the remaining three lacked annotations.

**Table 1.** Significantly associated genes that are common in TWAS-WW (FE and RE) and TWAS-WD (FE and RE). Only the genes with functional annotations available in AGPv4 assembly of maize genome are represented here. The column “Gene” refers to the Ensembl IDs of genes, the column “Chr” refers to the chromosomes on which these genes are located, and the column “Functions” represents the proteins they encode or the functions they perform.

| Gene | Chr | Functions |
| --- | --- | --- |
| Zm00001d034045 | 1 | Agamous-like MADS-box protein (AGL8) |
| Zm00001d035346 | 6 | Probable sucrose-phosphatase 2 (SPP2) |
| Zm00001d035377 | 6 | IQ domain-containing protein IQM3 |
| Zm00001d035390 | 6 | Resistance to sugarcane mosaic virus 1 *SCMV1* (*ZmTrxh*) |
| Zm00001d035409 | 6 | Absciscic stress-ripening protein 2 (ASR2) |

*Zm00001d034045,* or Agamous-like MADS-box protein (*AGL8*), is a transcription factor involved in meristem identity regulation (21). It is also known to control fruit development and floral transition in *Arabidopsis* (22). *Zm00001d035346* forms sucrose-phosphatase 2 (SPP2), which catalyzes the final step in sucrose biosynthesis (23). *Zm00001d035377* forms IQM3 protein, a calmodulin-binding protein involved in calcium signaling and is important for different development and growth-related traits (24). *Zm00001d035390* (*ZmTrxh*) is responsible for resistance to sugarcane mosaic virus in maize (SCMV) (25). This virus can cause a severe decrease in maize yield (26), which however was not the case in this MET. *Zm00001d035409* encodes abscisic stress-ripening protein 2 (*ASR2*), which is known for conferring drought resistance to different plant species such as Arabidopsis (27), chickpea (28), tomato (29), and rice (30). *ASR2* is also involved in resistance to salt stress in different species, such as durum and common wheat (31).

Comparing significantly associated genes between TWAS-WW and TWAS-WD, 23 genes were uniquely identified in WW condition, while 11 genes were uniquely identified in WD condition. Many of these significantly associated genes, in particular those unique to TWAS-WD, exhibited an important change in expression from WW to WD conditions (Fig. S2). Some notable genes include *Zm00001d007949*, responsible for zea apetala homolog1 (*ZAP1*), and *Zm00001d008939*, responsible for encoding *flowering locus T* (*FT*) protein. *ZAP1* is a developmental gene in maize that has also been reported to be expressed in maize under field drought stress (32). An increase in ear weights and kernel numbers in *ZAP1* knockdown mutants grown in different nitrogen regimes in the field has also been reported (33). These studies indicate an important role of *ZAP1* in maize grain yield. *FT* belongs to the *phosphatidylethanolamine-binding protein* (*PEBP*) family and is responsible for the activation of floral activating complex (FAC), which in turn upregulates more flowering genes such as *apetala-1* (*AP1*) and *leafy* (*LFY*) (34–36). *PEBP* has also been reported as a potential target for yield improvement in quinoa (36).

In all of the four cases of TWAS, i.e., TWAS-WW and TWAS-WD with FE and RE procedures, most of the genes found to be significantly associated with grain yield were present on chromosome 6, especially within the same region where strong associations were also observed with GWAS. This region was identified as carrying a large 2 Mbp presence absence variant (PAV) in previous studies (26,37–40).

### Transcriptomics Enhances the Resolution of Association Analysis

The region carrying the PAV polymorphism on chromosome 6 was observed to be strongly associated with grain yield (Fig. 1A, B). QTLs carrying large PAVs can cover wide regions because of important LD extent due to strong depletion of recombination within and around PAV, making it difficult to identify the causal gene. Moreover, performing association analysis on a PAV region can result in strong associations of all the SNPs present in the region with the trait, making it impossible to determine the causal gene(s). We hypothesize that TWAS can improve on the drawbacks of GWAS as it directly tests for associations between genes and trait, suffering less from LD (5). For this, TWAS and GWAS were performed on a reduced panel of 178 hybrids that all had the presence variant of this region. We discriminated lines carrying the PAV sequence by using Off Target Variant calling approach from Affymetrix Axiom. We tested only the SNPs and transcripts present in the region within the span of 20 Mb to 28 Mb on chromosome 6 (Fig. 2 and S3 for the RE and FE procedures, respectively). In the case of the RE procedure, the number of significant associations for the reduced panel was of two SNPs for GWAS, one gene for TWAS-WW, and one gene for TWAS-WD. The two significantly associated SNPs detected by GWAS were located near *Zm00001d035421* and *Zm00001d035429*. *Zm00001d035421* lacks annotation in AGPv4, while *Zm00001d035429* encodes for *Retrovirus-related Pol polyprotein LINE-1*. These two SNPs are in strong LD with other regions of the PAV, making it difficult to identify causal genes. In the case of TWAS, the *ZmTrxh* gene showed the strongest association with grain yield in both FE (Fig. S3) and RE (Fig. 2) procedures, while *ASR*2 was close to being significant. These genes are potential causal genes that could be targeted for further functional validations in future studies. In the case of the FE procedure, no significantly associated SNPs or transcripts were detected (Fig. S3).

**Fig. 2.**
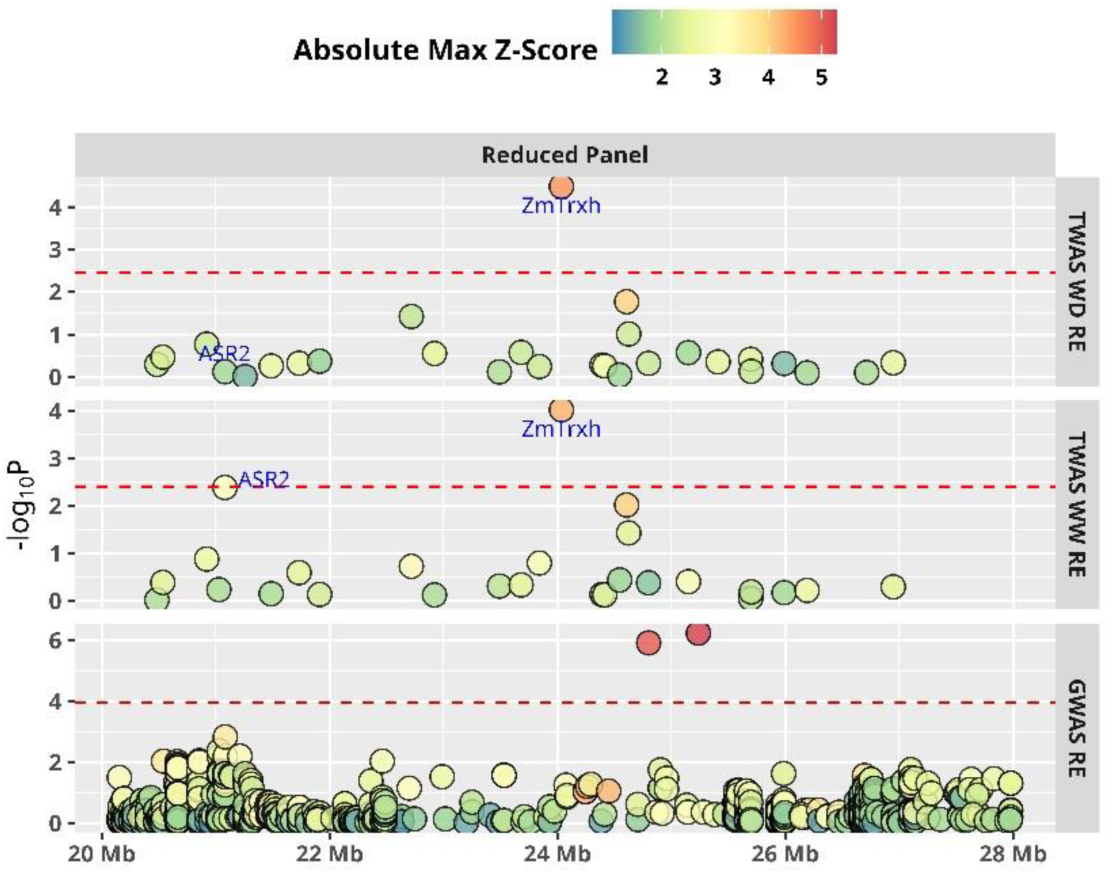
GWAS and TWAS on PAV region of chromosome 6 in the reduced genotype panel. The reduced panel is composed of the genotypes (178) having the presence variant. The meta-analysis was carried out based on the RE procedure. Each point represents a SNP in the case of GWAS and a transcript in the case of TWAS-WW and TWAS-WD. The color scale represents the absolute maximum Z-score of the corresponding SNP or transcript. The red dashed line represents the Bonferroni (10%) threshold. The X-axis represents positions in mega-base pairs (Mb), and the Y-axis represents the −log_10_ of the p-values.

### Effects of Transcripts on Grain Yield Depend on Environmental Conditions

The effects of transcripts on grain yield varied from one environment to another or even from one group of environments to another. For example, a transcript such as *ZmTrxh* had a stronger effect in water deficit and hot field trials compared to well-watered and cool trials (Fig. S4). Such trends become much clearer when the Z-scores are regressed against environmental covariates using the meta-regression test procedure (41). This procedure was used to identify if transcript effects variations correlated with environmental covariates. An FDR of 10% was applied in this procedure to control for type I errors. The results indicated that the effects of *ZmTrxh* and *ASR2* were significantly correlated with mean maximum temperature (Tmax.mean) and mean vapor pressure deficit (VPDmax.mean) (Fig. 3). These results indicate that the effects of these genes on yield were highly dependent on environmental factors and their effects were much stronger in stressful conditions.

**Fig. 3.**
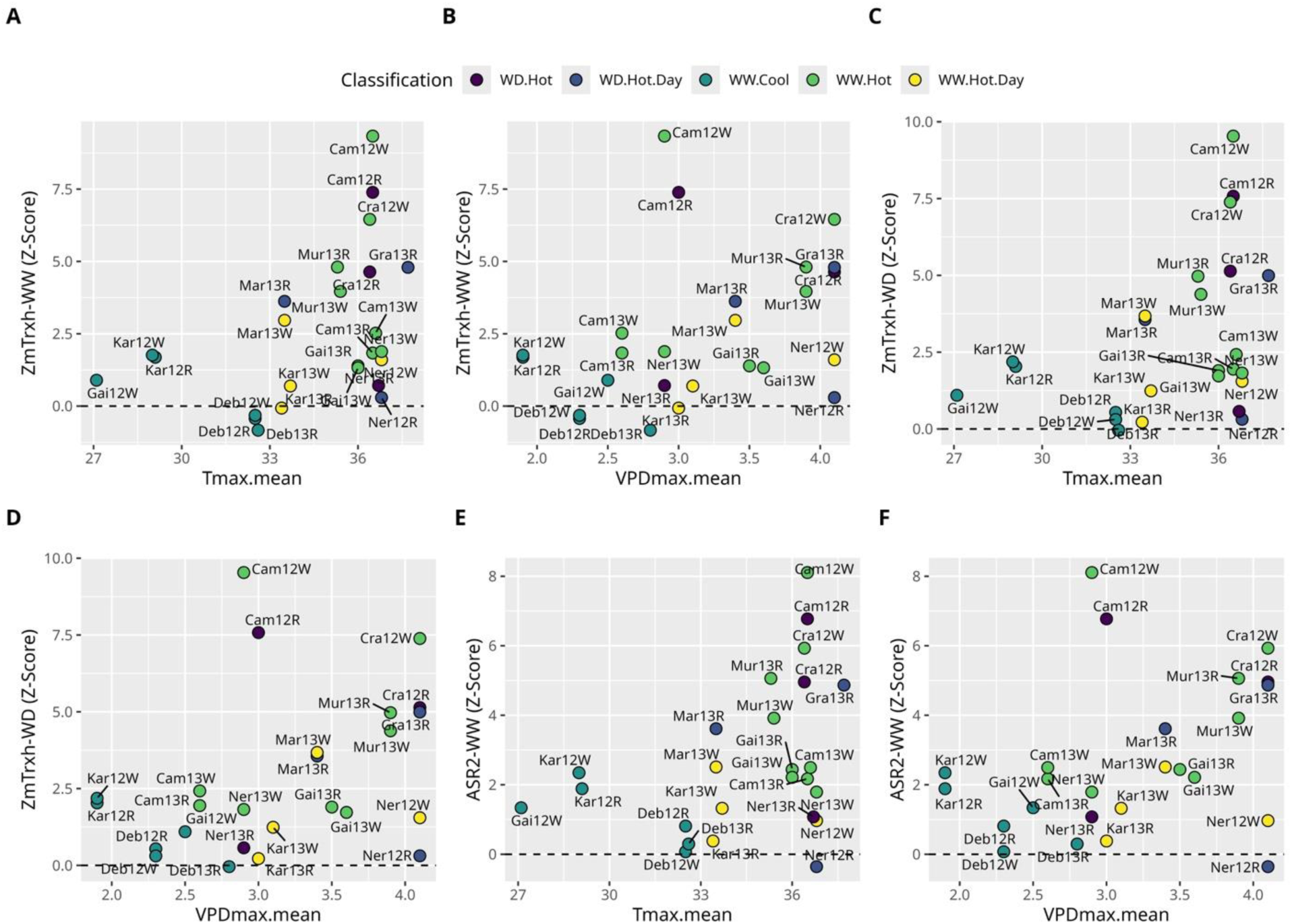
Comparison of the Z score of a given transcript within each environment against the value of the environmental covariates (ECs). The regression was carried out using the meta-regression procedure of De Walsche et al. (2025). Only the transcript and ECs that exhibited significant interaction are plotted. The X-axis represents the value of a given EC, and the Y-axis represents the Z scores of a given transcript in a given environment. The color scale represents different environmental classifications. The black dashed horizontal line represents zero. A) Z score of Zm00001d035390 (ZmTrxh) (WW) against Tmax.mean, B) Z score of Zm00001d035390 (ZmTrxh) (WW) against VPDmax.mean, C) Z score of Zm00001d035390 (ZmTrxh) (WD) against Tmax.mean, D) Z score of Zm00001d035390 (ZmTrxh) (WD) against Tmax.mean, E) Z score of Zm00001d035409 (ASR2) (WW) against Tmax.mean, and F) Z score of Zm00001d035409 (ASR2) (WW) against VPDmax.mean.

### Identification of potential epistatic relationship using eQTL analysis

Further on, we will refer to transcripts that are significantly associated with grain yield as biomarkers. Our objective here was to evaluate if the eQTLs of biomarkers were also associated to grain yield. We compared the *p-values* of SNPs from eQTLs analysis on biomarkers (in our case the associations were tested using a multi-locus approach to avoid redundancy between SNPs (42)) and the p-values of these same SNPs for their direct association with grain yield (GWAS) (43). The SNPs detected by eQTL analysis on the biomarkers were divided into cis and trans-eQTLs. Cis-eQTL SNPs were defined as all the significantly associated SNPs located at less than one megabase of the biomarker’s gene, while a trans-eQTL SNP was defined as all the significantly associated SNPs outside this window. In total across the 31 WW biomarkers, we found 69 significantly associated SNPs (eQTLs), with 25 as cis-eQTL SNPs and 44 as trans-eQTL SNPs (Table S4) using a multi-locus association model (42). Similarly, we found 58 significantly associated SNPs (eQTLs) across the 21 WD biomarkers, with 17 as cis-eQTL SNPs and 41 as trans-eQTL SNPs (Table S5).

The −log_10_ p-values of the cis and trans-eQTL SNPs were compared to the *p*-values of these same SNPs for their association with grain yield (standard GWAS). Overall Pearson correlations of 0.33 and 0.35 were observed between the *p-values* of SNPs for eQTL of WW biomarkers & GWAS-FE, and eQTL of WW biomarkers & GWAS-RE, respectively (Fig. 4A, B). Similarly, Pearson correlations of 0.53 and 0.55 were observed between the p-values of SNPs for eQTL of WD biomarkers & GWAS-FE, and eQTL of WD biomarkers & GWAS-RE, respectively (Fig. 4C, D). These correlations suggest that if the effect of an SNP is high on a biomarker, it is also high on grain yield. However, most of the SNPs significantly associated with a biomarker expression had an effect below the significance in the GWAS, likely due to a lack of power. Here, transcriptomics-based approaches helped increase detection power by detecting more SNPs associated to grain yield, directly or through regulation, potentially resulting in epistasis.

**Fig. 4.**
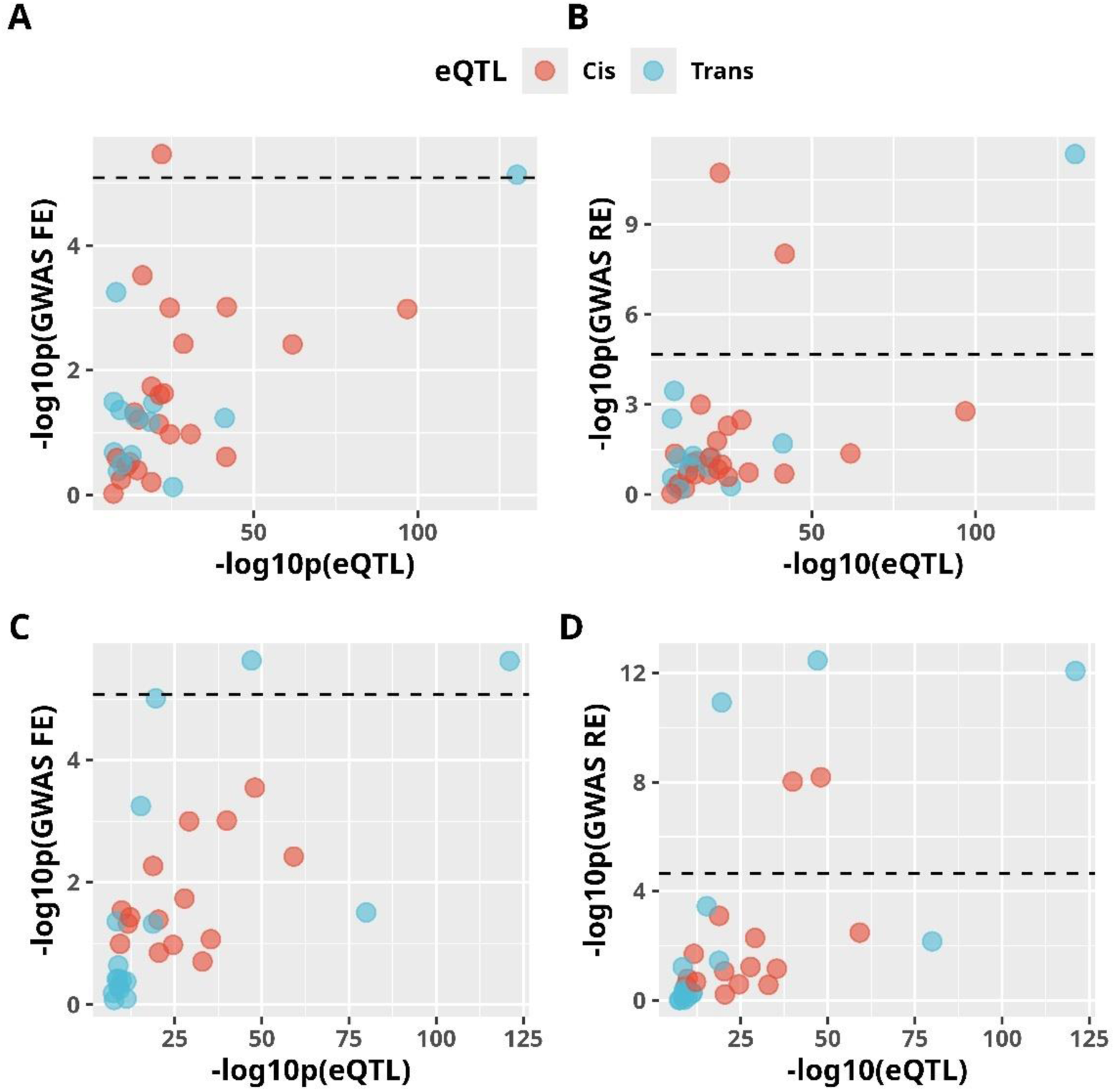
Comparison of the eQTLs SNPs for their p-values in eQTL analysis (horizontal axis) and GWAS (vertical axis). A) eQTLs of WW biomarkers versus GWAS-FE, B) eQTLs of WW biomarkers versus GWAS-RE, C) eQTLs of WD biomakers versus GWAS-FE, and D) eQTLs of WD biomarkers versus GWAS-RE. The red and blue colors represent cis-eQTL SNPs and trans-eQTL SNPs, respectively. The black dashed line represents the GWAS significance threshold as in Fig. 1A and B. Note that all the SNPs below the horizontal dashed line

### Important Contribution of Regulation to Genetic Variance and Genomic Prediction Accuracy

We evaluated the contribution of these cis- and trans-eQTLs to the genetic variance of grain yield in each trial using the following procedure (44): using a standard mixed model with a random genetic effect, we estimated the decrease of genetic variance when including a set of SNPs as fixed effects. The contribution of this set of SNPs to the genetic variance was estimated as the percentage of decrease of genetic variance. It appeared that the cis- and trans-eQTLs of biomarkers had an important contribution to genetic variance (Fig. 5, S5). Together they explained on average over the field trials 24,6% and 15,8% of the genetic variance for WW- and WD-eQTLs, respectively. The contribution of cis-eQTLs was higher (22,1% for WW) than the trans-eQTLs (14,1% for WW). Still the contribution of trans-eQTLs was high, indicating the potential important contribution of epistatic interactions to genetic variance. After dividing the contribution of the different sets of SNPs by the number of SNPs in the set, we observed a contribution to genetic variance of up to 1.79% per SNP in the case of cis SNPs identified from WW biomarkers (Fig. 6A), and up to 2.74% per SNP in the case of cis SNPs identified from WD biomarkers (Fig. 6B). The importance of cis SNPs identified from both transcript types was consistently higher in all environments compared to random samples of SNP (maximum contribution of 0.15%). A decrease of up to 0.88% per SNP was observed in the case of trans-eQTL SNPs identified from WW biomarkers (Fig. 6A), and 0.79% per SNP for those of WD biomarkers. Overall, it can be observed that cis-eQTL SNPs individually have a stronger effect than trans SNPs in explaining genetic variation, which is in accordance to the findings of several studies (45–47).

**Fig. 5.**
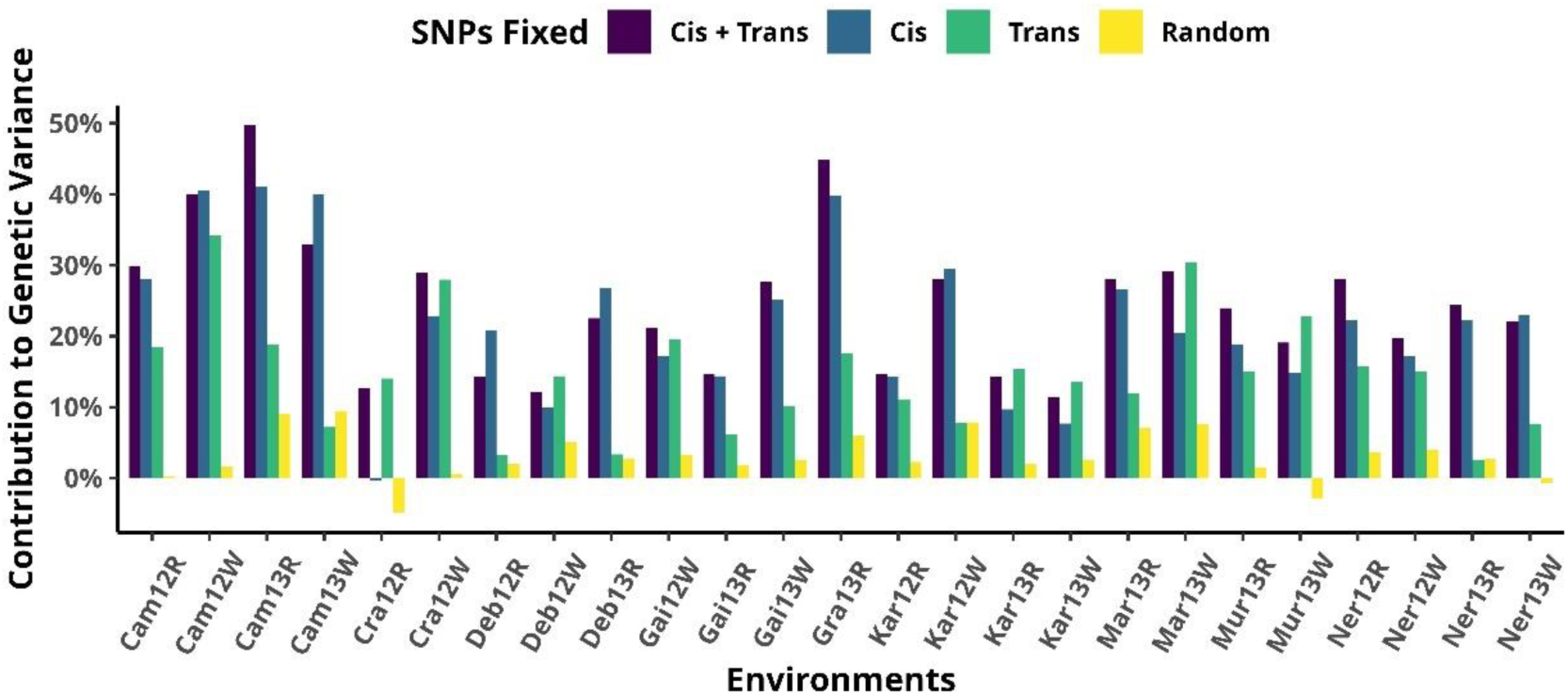
Contribution to genetic variance of the eQTL SNPs from the eQTL analysis on WW biomarkers. Cis: cis-eQTL SNPs, Trans: trans-eQTL SNPs, Random: random subsets of SNPs with as many SNPs as in Cis + Trans (the height of the bar corresponds to the average over 100 random samples).

**Fig. 6.**
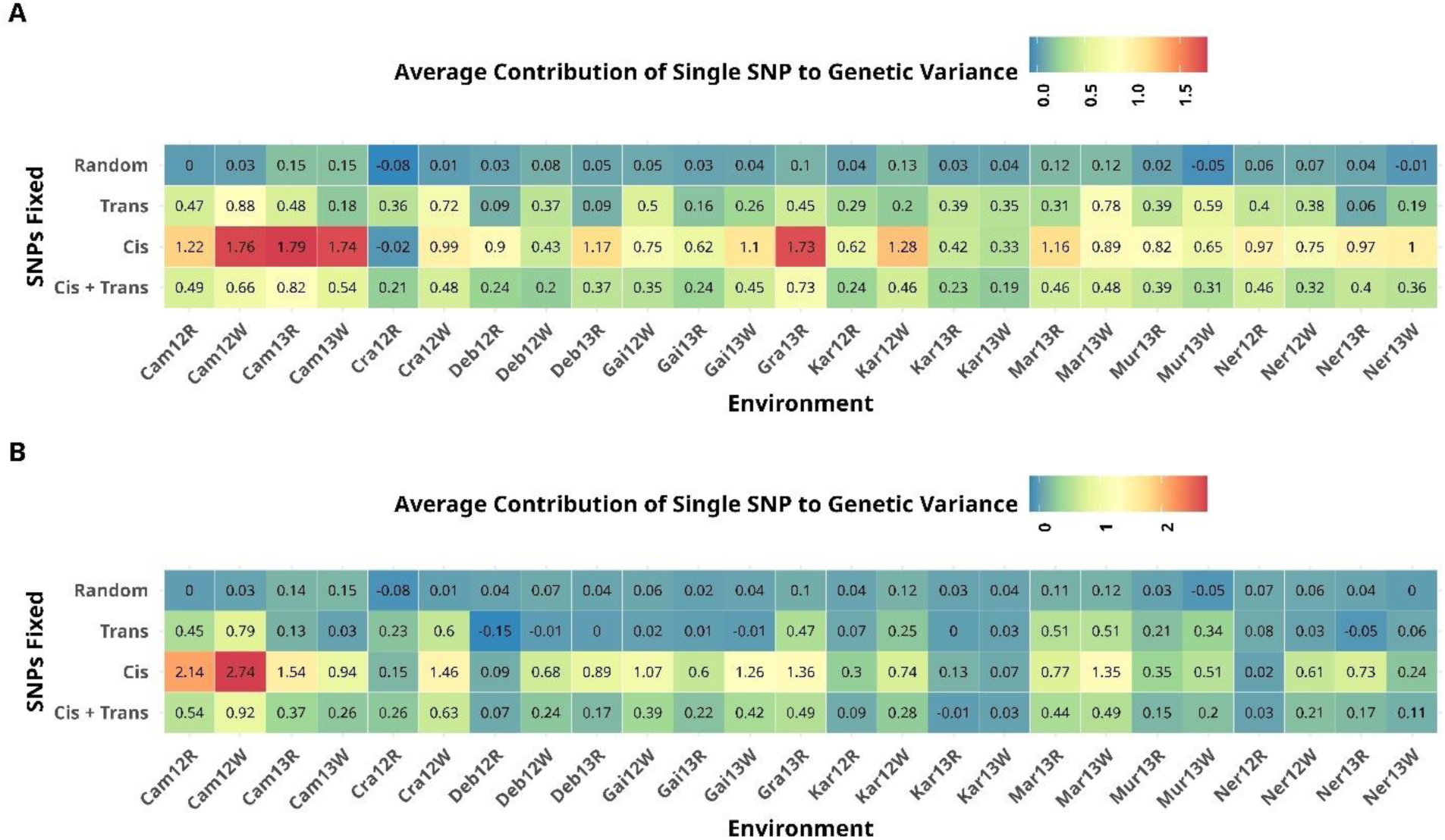
Contribution of cis and trans eQTL-SNPs to genetic variance. A) Percentage decrease of genetic variance per SNP identified from eQTLs of WW biomarkers, B) Percentage decrease of genetic variance per SNP identified from eQTLs of WD biomarkers. The contribution of eQTLs to genetic variance is estimated as the percentage of total genetic variance decrease when these eQTLs are included in the model as fixed effects.

Since these identified SNPs have a high contribution to genetic variance, we evaluated their efficiency to predict grain yield in comparison to random sets of SNPs of the same size. As it is necessary to keep the training and test sets separate in genomic prediction, we applied TWAS (FE and RE) and subsequent eQTL analyses only on the training set. Cis- and trans-eQTL SNPs identified in the training set were then used to make predictions in the test set. Results indicate that cis-eQTL SNPs consistently outperform both trans-eQTL SNPs and random sample (repeated 100 times) (Fig. 7A, B). Predictive ability (Pearson Correlation) of 0.43±0.11 and 0.39±0.12 across all the environments was observed in the case of cis-eQTL SNPs for WW and WD biomarkers, respectively (Table 2). These predictive abilities were much higher than what was obtained with random sets of SNPs (around 0.30), indicating the importance of the identified cis-eQTL SNPs. Unlike cis SNPs, trans-eQTL SNPs resulted in lower predictive abilities compared to the random sampling scenario.

**Fig. 7:**
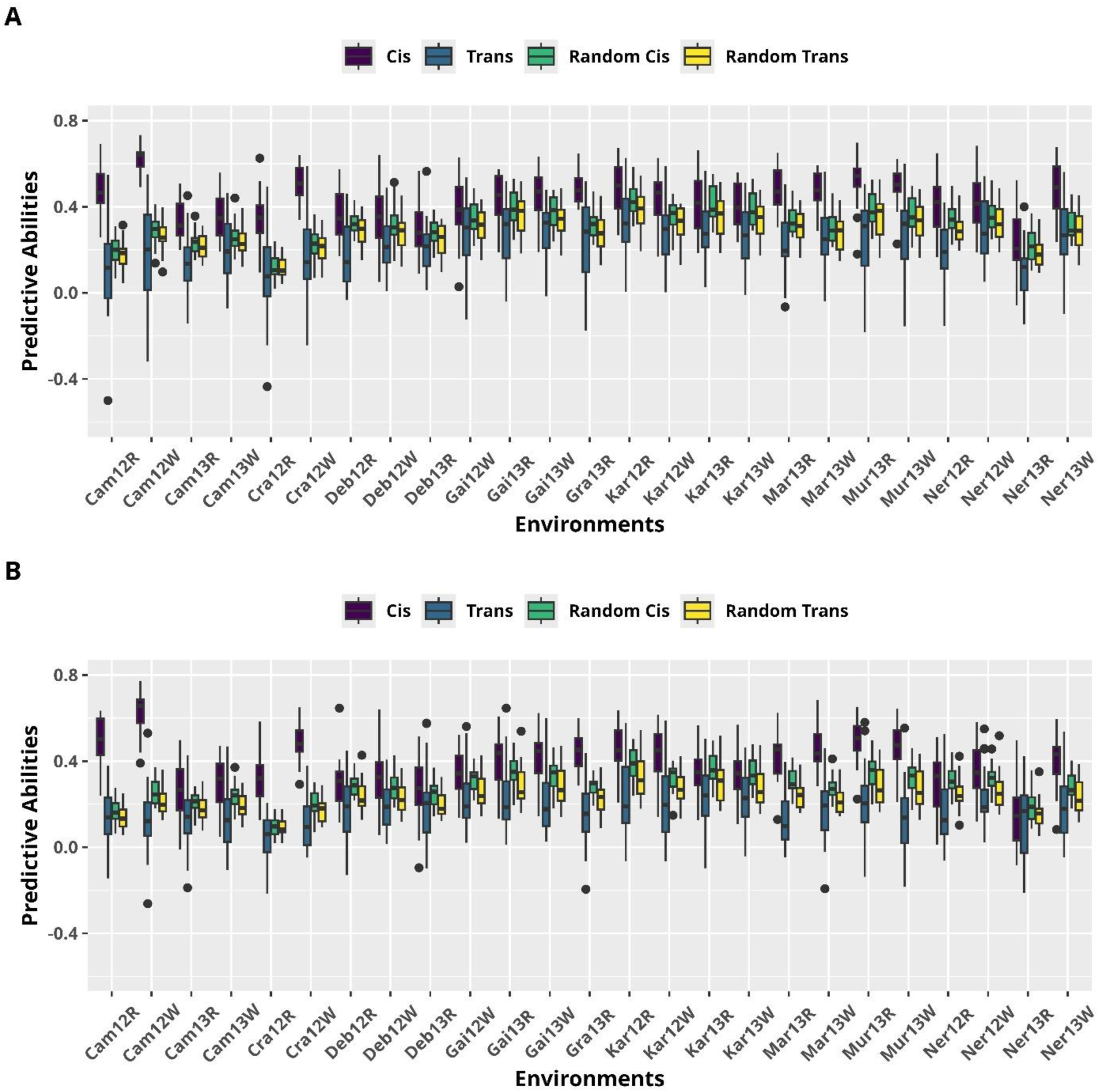
Predictive abilities of cis and trans eQTL-SNPs in comparison to random sets of SNPs. A) Predictive abilities of eQTLs of WW biomarkers or an equally sized random sample of SNPs, and B) Predictive abilities of eQTLs of WD biomarkers or an equally sized random sample of SNPs. Cis: cis-eQTL SNPs, Trans: trans-eQTL SNPs, Random Cis: random subsets of SNPs with as many SNPs as in Cis (100 independent random samples), Random Trans: random subsets of SNPs with as many SNPs as in Trans (100 independent random samples).

**Table 2.** Predictive abilities of prediction models averaged across all 25 trials and cross-validations.

| | Predictive Abilities (PA±SD) | |
| --- | --- | --- |
| SNPs in Prediction Model | WW biomarkers | WD biomarkers |
| Cis eQTLs | 0.43±0.11 | 0.39±0.12 |
| Trans eQTLs | 0.23±0.16 | 0.17±0.15 |
| Random (same size as Cis) | 0.31±0.07 | 0.28±0.06 |
| Random (same size as Trans) | 0.28±0.08 | 0.23±0.07 |

## Discussion

Maize grain yield is a complex trait likely influenced by many interacting genetic factors. This study strives to answer two key questions: How can we combine transcriptomics with standard GWAS approaches to improve our understanding of the genetic architecture? And is this combination able to increase detection power and resolution, when omics data are collected from a controlled condition to model the genetic architecture of a complex trait observed in a field setting? This study utilizes a comprehensive dataset of maize grain yield of 244 genotypes in a multi-environmental trial conducted in contrasted watering and thermic conditions across Europe and an external site in Chile (15). Transcriptomic profiles were obtained for the same panel grown on a platform in well-watered (WW) and water deficit (WD) conditions. This dataset was analyzed with an established association study model (18) applied to each field trial, followed by a recently developed meta-analysis approach (41). One novelty of our study was to apply the meta-analysis to the omics data, taking into account the contrasted environmental conditions and GxE interactions. And most importantly, GWAS, TWAS and eQTL analyses were combined to detect regulatory regions involved in the establishment of grain yield. Resultantly, key QTL regions, genes and cis- and trans-regulatory regions that affect grain yield in an environment conditions-dependent manner were identified.

For GWAS, the FE meta-analysis procedure revealed significantly associated regions on chromosomes 3 and 6, while the RE procedure revealed different regions on chromosomes 1, 2, 3, 6, 8, and 10. QTLs for grain yield in similar regions were also identified in past (48) and also on the same material but with much less molecular markers (41). We were able to associate the regions identified via the FE procedure to 15 unique genes and the ones from the RE procedure to 41 unique genes by considering a 10 kb window upstream and downstream of all significantly associated SNP positions. Most of these genes either lacked a functional annotation or were pseudogenes. Hence, further studies are required for the functional validation of these genes. A GO analysis via agriGO V2.0 identified 35 of these genes as protein-coding (most lacking functional validation) and identified an enrichment genes (5 genes) related to ubiquitin-protein transferase activity, which are known to be associated with abiotic stress response in maize (20). The same molecular function was also validated to be involved in abiotic stress in two bread wheat varieties (49).

In the case of TWAS, eight genes were identified as common across both watering conditions (WW and WD) and meta-analysis approaches (FE and RE). These genes could not be associated with a particular GO term via agriGO V2.0, likely because some lacked functional information and others belonged to different functional domains or pathways. AGPv4 genome assembly gives functional information for 5 of these genes (Table 1). We could link these functions to maize flowering, growth and development, and resistance to biotic and abiotic stresses (21–26,28). Moreover, 23 genes were identified with the WW transcripts but not with the WD transcripts, while 11 were with the WD transcripts but not with the WW transcripts. We also observed clear differences between the expressions in the two conditions for these genes.

The region on chromosome 6 indicated a strong association with grain yield in both GWAS and TWAS. This region was shown to include a PAV (37,38), including numerous genes with nearby SNPs in strong LD, making it difficult to identify the actual causal gene. Transcripts are less likely to be affected by LD in association analysis (5). This property can be useful in identifying a causal gene, as shown using TWAS on the reduced panel (the set of genotypes having the presence). TWAS can be much more useful in this case, as it reveals the association of potential causal genes inside the PAV because of variation in the expression levels of each gene of the segment. The gene *ZmTrxh* exhibited the strongest association, followed by *ASR2* in the reduced panel, indicating that the expression of these two genes affects yield. These results indicate the importance of transcriptomic or expression data in identifying the potential causal genes (50). In line with the findings of this study, transcriptomic data have been useful in unraveling the genetics of plant architecture, disease resistance, and flowering in previous studies (38,51–53). However, in most of these studies, omics data were collected in the same conditions under which phenotypes were measured. We demonstrated that platform omics can also provide insights into the genetic determinants of field traits.

Another important result to note here is that the effects of the transcripts can vary from one environment to another depending on the underlying climatic conditions. For example, a significantly associated gene set on chromosome 6 strongly affected grain yield in stressed environments (Fig. S4). However, their effects were much smaller in the case of favorable environments (classified as the “WW.Cool” environmental group). These results are in line with the fact that these genes have already been identified in different studies to be responsible for response to stresses (25,27,31,32). Meta-regression further revealed that the effects of two genes, *ZmTrxh* and *ASR2,* are significantly dependent on different environmental covariates, including Tmax.mean and VPDmax.mean (Fig. 3). Their effects were particularly high in the environments in which both days and nights were hotter, indicating a strong dependence on heat stress.

An important drawback of TWAS is that only association with genes are considered, meaning that all the regulatory regions outside genes remain unexplored. But we showed that eQTL analyses of the transcript associated with yield were able to detect cis and trans regions outside genes. Most of these regions were undetected by GWAS (Fig. 4), meaning that the combination of TWAS and eQTL analyses can significantly enhance detection power. The use of transcripts appears particularly interesting because they have a much simpler genetic determinism compared to complex traits. Additionally, since they are far less numerous than SNPs, they help reduce the multiple testing burden. Such an approach is quite advantageous as it can highlight SNPs that play regulatory roles potentially leading to epistatic interactions, such as the SNPs present in trans-eQTLs of a transcript related to grain yield. Upon further exploring the importance of these cis and trans-eQTLs, we found that both greatly contribute to the genetic variance of grain yield. Between the two, cis-eQTLs tended to explain globally and individually a higher per SNP variance. One explanation could be that we have much less power to detect the trans-eQTLs as their effects are smaller than the cis, meaning that some trans regions still remain unidentified. The cis-eQTL SNPs were also particularly useful in genomic predictions, where they consistently performed much better than random samples of SNPs of the same size, with a gain of 39% (cis-eQTLs of WD biomarkers) to 52% (cis-eQTLs of WW biomarkers) in predictive ability in comparison to random sets of SNPs (Fig. 7, table 2). This information is particularly useful to design bio-informed genomic prediction models (54–57), able to predict more accurately, in particular genetically distant material (58,59). On the contrary, the trans-eQTLs did not increase predictive ability in comparison to random SNPs. One explanation to this difference could be that the trans regulation contributes more than cis regulation to the response to the growth conditions with rapid and reversible changes (60,61), meaning that they are probably more affected than the cis by the important environmental distances between the platform and the fields.

In our study, we demonstrated that transcriptomic data obtained from controlled conditions of a platform can be used to some extend to dissect the genetic architecture of field productivity traits. This result was expected, as it was shown in numerous studies that omics from controlled conditions could be used to predict efficiently field traits (10,11). However, predicting and modelling genetic architecture are two different objectives, the latter being much more ambitious. For instance, we can suppose that omics are able to capture genetic relatedness, which is a key element of genomic prediction (62,63), whereas it seems more challenging to relate the expression of a gene on a platform and the elaboration of yield in the field. One explanation of why it worked in our study, is that there are certainly common biological processes regarding the adaptation to the growth conditions between the platform and the fields because of some common stressing factors (e.g. water stress). In any case, this opens the way to promising applications in the field of plant genetics and breeding, as omics cannot yet be applied systematically to all field trials because of the high costs.

## Conclusions

Because of the cost for omics acquisition, it is still not an option to measure them systematically in a multi-environment trial. This is the reason why in most studies, the acquisition is done once and for all in one controlled condition, often an indoor platform (10,64). Our study is the first, to our knowledge, to demonstrate that this kind of experiment is efficient to increase our understanding of the genetic architecture of a trait measured in a multi-environment trial. Overall, the results indicate that an integrative approach combining transcriptomics and genomics can improve resolution and detection power in particular for regulatory regions. Our study opens the way to the routine use of integrative multi-omics approach in crop genetics.

## Methods

### Plant Material and Genotyping

A hybrid panel of 244 maize dent lines crossed to a single flint parent (UH007) was used in this study (15). The flowering time of the panel was restricted to a window of seven days. The panel traced back to three heterotic groups: Iodent, Lancaster, and Stiff Stalk. The panel was genotyped using a 50K Infinium HD Illumina array (65), a 600K Axiom Affymetrix array (66), and genotype by sequencing (GBS) (48), resulting in 978,134 SNP markers for all the genotypes. We filtered out SNPs with minor allele frequencies (MAF) lower than 0.05 to remove fixed or nearly fixed variants. Eventually, we worked with a set of 747,262 SNPs for the association studies.

### Phenotyping Data

The hybrid panel of 244 genotypes was evaluated in the DROPS (https://cordis.europa.eu/project/id/244374) project (15) for grain yield and related traits in 25 different trials under irrigated and rainfed conditions along western to eastern Europe and an external site in Chile. Genotypic means for grain yield were estimated using a mixed model with genotypes and replicates as fixed effects and spatially correlated errors as random effects (15). The model was fitted using ASReml-R (67,68), and the best linear unbiased estimators (BLUEs) of the genetic effects were used in this study. Different environmental covariates, including soil water potential, maximum temperature, night temperature, and vapor pressure deficit, were also measured during DROPS trials (15). Another covariate, radiation intercepted, was estimated through simulations using the APSIM crop growth model (69,70) and included in the environmental covariates (70). The mean value of these covariates for each trial was computed across three developmental phases of plants, i.e., early development, flowering, and grain-filling. These covariates have been used to classify trials into five classes (70): water-deficit hot (*WD Hot*), water-deficit hot day (*WD Hot Day*), well-watered hot (*WW Hot*), well-watered hot day (*WW Hot Day*), and well-watered cool (*WW Cool*).

### 3’ RNA seq Gene Expression

In the Spring 2013 platform experiment, 3’ RNA seq gene expression analysis was performed on mature leaf tissue samples of 244 genotypes for both WW and WD treatments (11). Each genotype had two replicates per condition, resulting in four leaf samples (one per plant). The new generation sequencing (NGS) libraries were synthesized using the QuantSeq-3’ mRNA-Seq Library Prep Kit FWD (Lexogen, Vienna). According to the manufacturer’s recommendations, the libraries were sequenced on a Next Seq 550 (Illumina, San Diego, USA) using a High-output flow cell. The resulting sequences were aligned to genome version 4 of B73 maize. Only reads aligning to a single gene and with at least one count per million reads in at least 10% of the samples were considered, which resulted in 20,642 transcripts for the WD treatment and 20,475 transcripts for the WW treatment. For each transcript, marginal genetic means were computed using a linear model, including the sampling hour, repetition, and genotypes as fixed effects and rows and columns as random effects. The same model with genotypes as random effects was also fitted to compute broad sense heritabilities. Ali et al. contains further details on transcript extraction, quantification, correction, and normalization (11). For the analyses in this study, transcripts with heritability lower than 0.4 or with a variance close to zero were removed, resulting in 6,980 WW transcripts and 6,997 WD transcripts.

### Statistical Models and Meta-Analyses on the Full Panel

We fitted a standard mixed model for association mapping accounting for relatedness (18) using the MM4LMM R package (71).

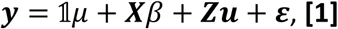

where ***y*** is the vector of adjusted phenotypes, 𝟙 is a vector of 1s, *μ* is the overall mean, ***X*** is the vector of genotype values of a given SNP in the case of GWAS, or the vector of pre-treated expressions of a given transcript in the case of TWAS. *β* is the corresponding fixed effect. ***Z*** is the design matrix for the random vector of genetic effects ***u***, modeled as 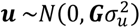, with ***G*** being the genomic relationship matrix obtained from genome-wide SNPs by following the VanRaden approach (72). 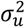 is the random genetic variance. ***ε*** corresponds to the vector of random errors modeled as 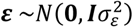, with ***I*** the identity matrix and 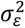 the random error variance. GWAS and TWAS models (with WW and WD transcripts) were applied to each grain yield trial independently.

After obtaining the *p-values* of the GWAS and TWAS in each trial, we applied the meta-analysis approaches “metaGE” (41) including the fixed and random effects procedures. The fixed effect procedure tests if the SNP or transcript effect is constant for all the environments or at least within prespecified groups of environments, while the random effect procedure tests for both constant and variable effects in the different environments. Further details of the procedures can be found in De Walsche (41). The *p-*values of the meta-analyses were corrected using a 10% false discovery rate (*p_FDR_*). Any SNP or transcript with a *p_FDR_*<0.10 was considered significantly associated with the trait.

### Statistical Models and Meta-Analyses on the Reduced Panel

Millet (15) identified a set of 19 SNPs originating from the 600K SNP panel (66) belonging to a PAV on chromosome 6. Based on the presence of these SNPs, 178 genotypes were selected for having the presence variation. These genotypes will be referred to as the reduced panel, on which we also applied GWAS and TWAS. SNPs or transcripts with *p*-values higher than 0.6 (default value proposed by the metaGE coauthors) were used to estimate genetic correlations between different environments. After following FE and RE procedures of meta-analysis on the reduced panel, only SNPs and genes present in the genomic region between 21Mb to 28Mb (corresponding to the PAV) were tested for association with yield. As the number of SNPs or transcripts was smaller in this case, Bonferroni (10%) correction was applied to correct for multiple testing.

### Meta-regression Test

De Walsche (41) introduced a meta-regression procedure to identify QTLs whose effect variations correlate with environmental covariates. This procedure can also be used for transcriptomic data. In this case, it tests the Z-scores of each transcript from all the environments as a function of quantitative environmental covariates. This test was applied to both WD and WW transcripts independently. An FDR of 10% was also applied to correct for multiple testing.

### eQTL analysis of the biomarkers to detect cis and trans regulation

We applied eQTL analyses to the transcripts significantly associated with grain yield (biomarkers) (43). It implies that if an SNP affects a transcript, and that transcript affects a trait, then the SNP should influence both the transcript and the trait. For the eQTL analyses, we applied the multi-locus mixed-model (MLMM) strategy (42). MLMM extends standard GWAS methods by incorporating both forward and backward elimination of SNP cofactors within a linear mixed model framework. This approach accounts for population structure and polygenic background by iteratively re-estimating genetic and residual variances at each step. To select the final model, we applied the multiple-Bonferroni criterion (mBonf) (at 10%) as described in the original study. This method allowed us to identify multiple independent associations while controlling for false discoveries and confounding due to relatedness. eQTL analysis was only applied to the transcripts that were found to be significantly associated with grain yield. The chromosomal location of the underlying gene was used to distinguish between cis and trans eQTLs. SNPs at less than one megabase from the biomarker’s gene were regarded as cis-eQTL SNPs and the ones at more than one megabase or present on a different chromosome were regarded as trans-eQTL SNPs.

### Contribution of eQTL SNPs to Genetic Variance and Genomic Prediction Accuracy

To evaluate the contribution of eQTL SNPs to grain yield, we performed both variance component analysis and genomic prediction using sets of significantly associated eQTL SNPs. For variance estimation, we used four scenarios: (1) Cis+Trans eQTLs, (2) Cis eQTLs only, (3) Trans eQTLs only, and (4) Random SNP sets sampled 100 times, each with the same number of SNPs as the Cis+Trans set. We obtained principal components (PCs) from genomic relationship matrix (15,44) from each of the four above mentioned sets of SNPs. Only the PCs following the Kaiser Criterion were included in the following linear model:

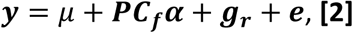

where ***y*** is the vector of adjusted means of grain yield, *μ* is the overall mean. *PC_f_* represents the selected PCs, and **α** their effects, ***ɡ_r_*** is the vector of random genetic effects that are considered to follow a normal distribution 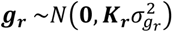. Here, ***K_r_*** represents the genomic relationship matrix computed based on the remaining SNPs. ***e*** represents the vector of random errors, i.e., 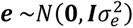. For each environment and each set of SNPs, we compared the genetic variance estimate 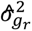 when the PCs on the SNP sets were included or excluded from the model. The contribution of a set of SNPs to genetic variance was estimated as its decrease when the corresponding PCs were included.

In the case of genomic predictions, four scenarios were considered: (1) Cis eQTLs, (2) Trans eQTLs, (3) Random Cis, and (4) Random Trans. In both Random Cis and Random Trans (random sets of SNPs with the same number of SNPs as in the Cis eQTL and Trans eQTL sets, respectively), the random sampling was repeated 100 times. We applied a 5 folds 5 repeats cross validation scheme, with one fold as test set and four folds as training set, and to avoid any biasness, all the analyses (TWAS, meta-analysis, and eQTL analyses) were performed only on the training set. The SNPs identified as eQTLs in the training set were used to make predictions in the test set. The following prediction model was fitted in all four scenarios:

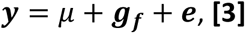

where ***y*** and *μ*, and ***e*** are the same as defined in equation 2. ***ɡ_f_*** represents the vector of random genetic effects mediated through selected feature (Cis, Trans, Random Cis, and Random Trans), with the following assumption: 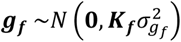. Here ***K_f_*** is the genomic relationship matrix computed from selected feature SNPs. Like in the case of variance estimation, the four scenarios of genomic prediction were also applied independently to each environment.

## Declarations

### Data Availability

Field grain yield and environmental covariates can be found at: https://doi.org/10.1104/pp.16.00621.

The genotyping data can be found at: https://doi.org/10.15454/AEC4BN.

The transcriptomics data can be found at: https://doi.org/10.57745/4NGRWR.

### Ethics approval and consent to participate

Not applicable

## Acknowledgements

We are thankful to all the DROPS and Amaizing partners for the discussions and the production of the datasets. We thank François Chaumont, Clémentine Vitte, Laurie Maistriaux, and Hervé Duborjal for the planification, the acquisition and acquisition of the transcriptomic data. We thank Hervé Duborjal for the first treatments of the transcriptomic data, and Maxime Laurent and Bertrand Huguenin-Bizot for the computation of the adjusted means and heritabilities.

## Authors’contributions

AC, SN, MBN designed the field and/or platform experiments. BA analyzed the data under the supervision of RR, AC, TMH, SN, MBN and LM. BA and RR wrote the manuscript. The study was supervised by RR. All authors have reviewed the manuscript, read and approved the final version.

