## Supplementary information for "Unravelling the Genetic Architecture of Field Traits through Multi-Omics Platform Data integration"

### Supplementary Figures

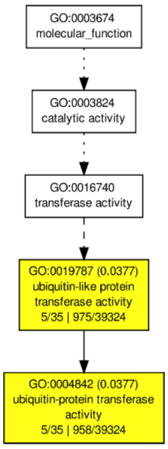


*Fig. S1. Gene Ontology (GO) enrichment analysis. GO analysis was performed via AgriGO v2 on genes that exist in close proximity to SNPs that are significantly associated with grain yield. A window of 10 kb upstream and downstream of SNP positions was used to identify the nearest gene. The diagram is a step-by-step hierarchy of parent and child terms. The numbers within the parenthesis represent the p-values indicating the association of the genes with the given GO term.*

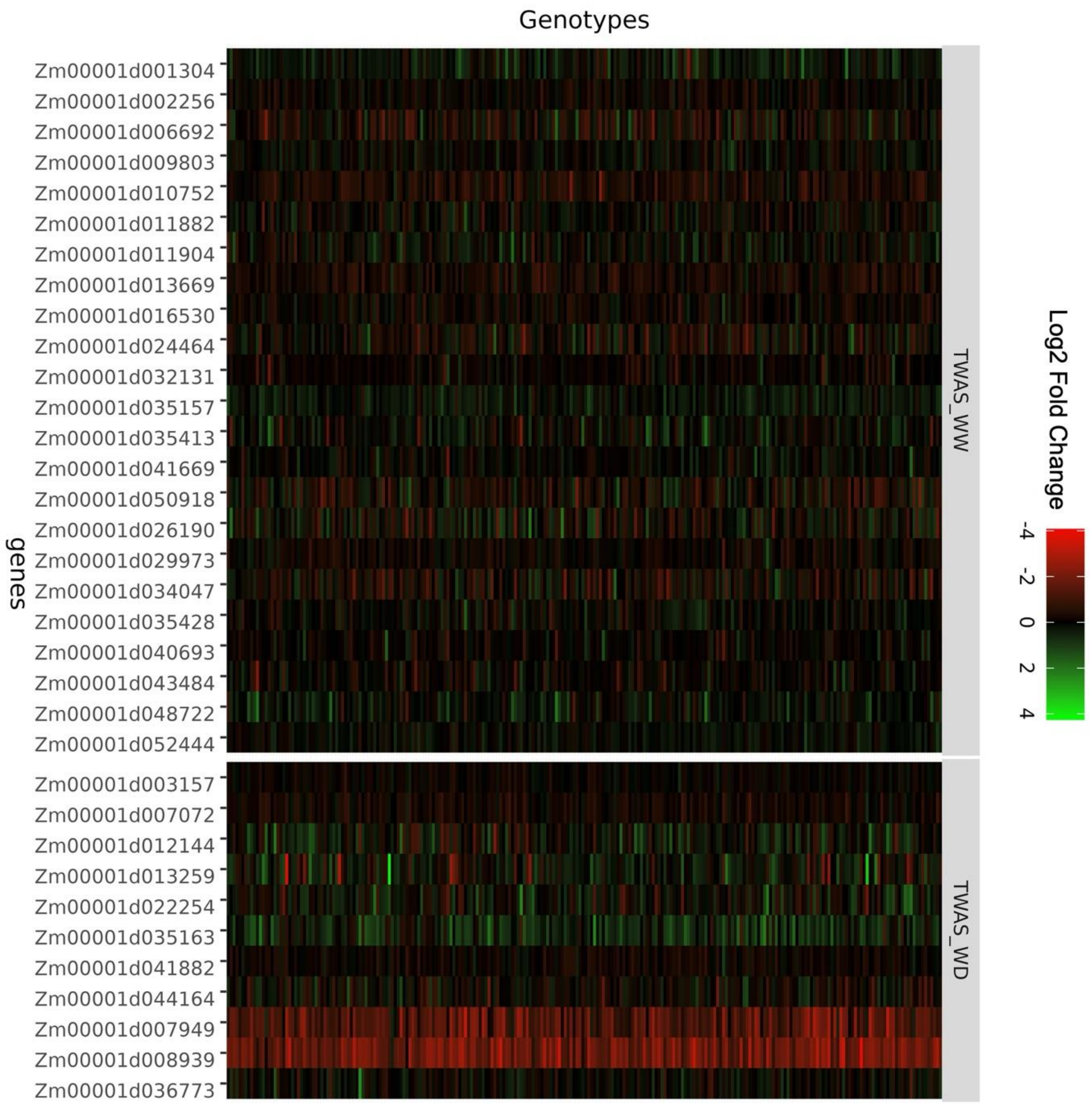


*Fig. S2. Differential expression of genes unique to TWAS-WW and TWAS-WD. The differential expression is taken as the log2 transformed expressions of $T_{WW}$ minus $T_{WD}$ ($T_{DIF} = T_{WW} - T_{WD}$). $T_{WW}$ and $T_{WD}$ being the count per million of the transcript in WW and WD conditions, respectively. The X-axis represents different genes. The Y-axis represents different genotypes. Each box represents $T_{DIF}$ for a given gene and genotype. The color represents a fold change in expression. The facet on the left represents significant genes unique to TWAS-WW, and the right one represents significant genes unique to TWAS-WD.*

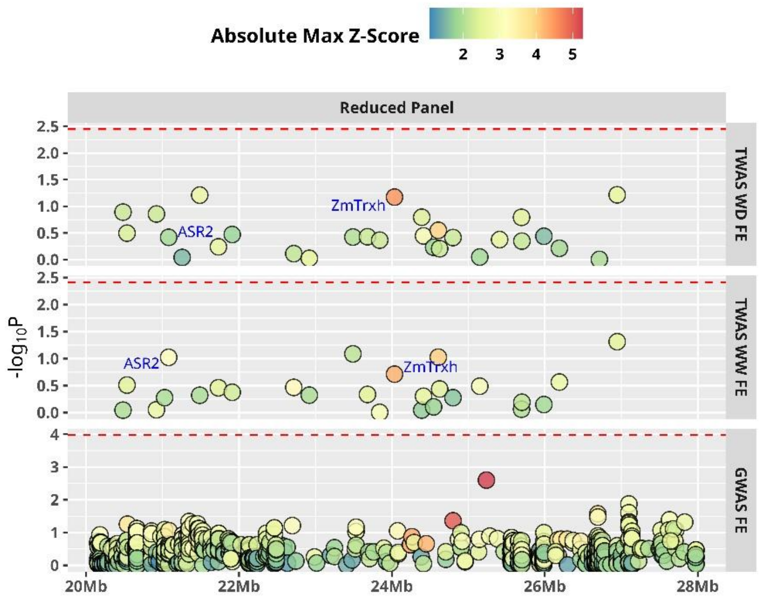


*Fig. S3. GWAS and TWAS on PAV region of chromosome 6 in the reduced genotype panel. The reduced panel is composed of the genotypes (178) having the presence. The meta-analysis was carried out based on the FE procedure. Each point represents a SNP in the case of GWAS and a transcript in the case of TWAS-WW and TWAS-WD. The color scale represents the absolute maximum effect of the corresponding SNP or transcript. The red dashed line represents the Bonferroni (10%) threshold. The X-axis represents positions in mega-base pairs (Mb), and the Y-axis represents the -log$_{10}$ of the p-values.*

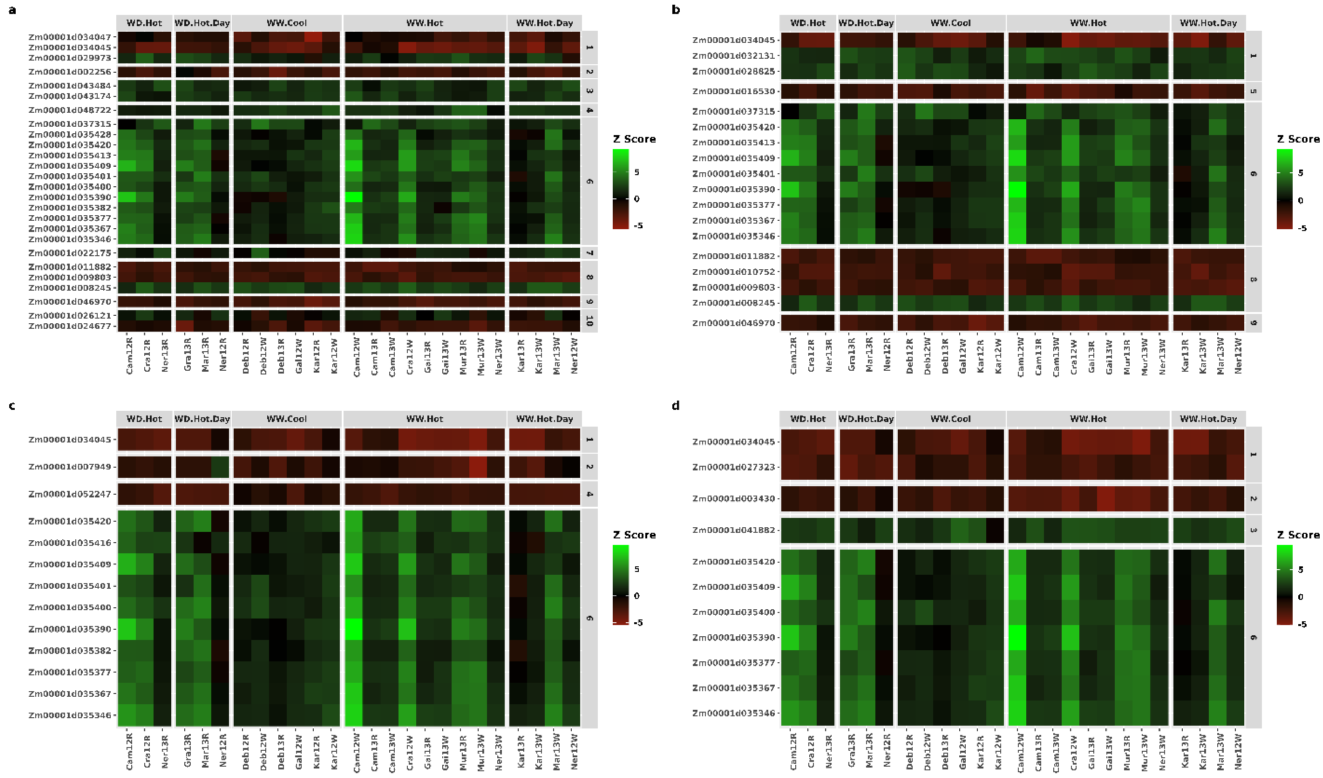


*Fig.S4. Comparison of Z scores of transcripts within each environmental classification. The X-axis represents different grain yield trials clustered into environmental scenarios, and the Y-axis represents genes found to be significant for a given version of TWAS: A) TWAS-WW-RE, B) TWAS-WW-FE, C) TWAS-WD-RE and D) TWAS-WD-FE. The color scale represents Z scores.*

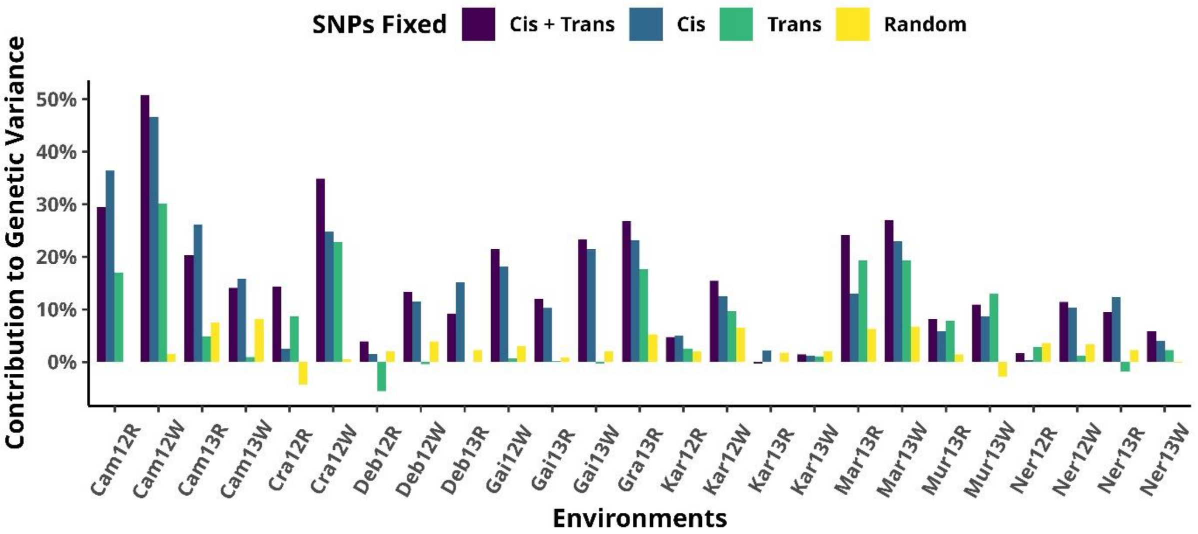


*Fig. S5. Contribution to genetic variance of the eQTL SNPs from the eQTL analysis on WD biomarkers. Cis: cis-eQTL SNPs, Trans: trans-eQTL SNPs, Random: random subsets of SNPs with as many SNPs as in Cis + Trans (the height of the bar corresponds to the average over 100 random samples).*

*Table S1: Genes identified using a 10kb window upstream and downstream of significant SNPs identified via GWAS with Fixed Effect (FE) procedure. "Gene" corresponds to Ensembl gene IDs, "chr" indicates the chromosome number, "Start" and "end" represent gene coordinates, "strand" represents positive or negative strand, and "description" represents gene functions.*

| gene | chr | start | end | strand | description |
|---|---|---|---|---|---|
| Zm00001d042256 | 3 | 1.58E+08 | 1.58E+08 | + | Protein kinase superfamily protein |
| Zm00001d042257 | 3 | 1.58E+08 | 1.58E+08 | - | Photosynthetic NDH subunit of subcomplex B 5 chloroplastic |
| Zm00001d042268 | 3 | 1.58E+08 | 1.58E+08 | - | Receptor-like protein kinase 5 |
| Zm00001d035345 | 6 | 22712369 | 22715419 | + | Retrovirus-related Pol polyprotein LINE-1 |
| Zm00001d035367 | 6 | 23489630 | 23490985 | + | Zm00001d035367 |
| Zm00001d035371 | 6 | 23595546 | 23596578 | + | Zm00001d035371 |
| Zm00001d035373 | 6 | 23598971 | 23600643 | + | Zm00001d035373 |
| Zm00001d001313 | 6 | 23743609 | 23746529 | - | Zm00001d001313 |
| Zm00001d035429 | 6 | 25236764 | 25238668 | + | Retrovirus-related Pol polyprotein LINE-1 |
| Zm00001d035240 | 6 | 16276743 | 16279273 | + | Zm00001d035240 |
| Zm00001d035232 | 6 | 15811792 | 15812517 | + | Zm00001d035232 |
| Zm00001d035227 | 6 | 15529000 | 15529392 | + | Zm00001d035227 |
| Zm00001d035228 | 6 | 15529973 | 15536315 | - | Methyl-CpG-binding domain protein 4-like protein |

| | | | | | |
|---|---|---|---|---|---|
| Zm00001d035229 | 6 | 15530206 | 15536423 | - | Methyl-CpG-binding domain protein 4-like protein |
| Zm00001d035224 | 6 | 14927596 | 14930694 | - | protein_coding |

*Table S2: Genes identified using a 10kb window upstream and downstream of significant SNPs identified via GWAS with Random Effect (RE) procedure. "Gene" corresponds to Ensembl gene IDs, "chr" indicates the chromosome number, "Start" and "end" represent gene coordinates, "strand" represents positive or negative strand, and "description" represents gene functions.*

| gene | chr | start | end | strand | description |
|---|---|---|---|---|---|
| Zm00001d034173 | 1 | 2.86E+08 | 2.86E+08 | - | Nuclear pore complex protein NUP58 |
| ENSRNA049468892 | 1 | 2.86E+08 | 2.86E+08 | - | snRNA |
| Zm00001d034174 | 1 | 2.86E+08 | 2.86E+08 | - | Zm00001d034174 |
| Zm00001d034175 | 1 | 2.86E+08 | 2.86E+08 | - | CCG-binding protein 1 |
| Zm00001d040537 | 3 | 48718848 | 48719919 | + | Zm00001d040537 |
| Zm00001d040540 | 3 | 48724704 | 48725590 | - | Zm00001d040540 |
| Zm00001d035149 | 6 | 7048494 | 7050657 | - | Putative endo-1%252C4-beta-mannosidase family protein |
| Zm00001d001296 | 6 | 7051470 | 7051729 | + | Zm00001d001296 |
| Zm00001d035150 | 6 | 7051761 | 7051964 | + | Zm00001d035150 |
| Zm00001d035151 | 6 | 7062012 | 7065762 | - | Zm00001d035151 |
| Zm00001d035345 | 6 | 22712369 | 22715419 | + | Retrovirus-related Pol polyprotein LINE-1 |
| Zm00001d035367 | 6 | 23489630 | 23490985 | + | Zm00001d035367 |
| Zm00001d035371 | 6 | 23595546 | 23596578 | + | Zm00001d035371 |
| Zm00001d035373 | 6 | 23598971 | 23600643 | + | Zm00001d035373 |
| Zm00001d001313 | 6 | 23743609 | 23746529 | - | Zm00001d001313 |
| Zm00001d035393 | 6 | 24064700 | 24069515 | + | Sucrose synthase 3 |
| Zm00001d035396 | 6 | 24239055 | 24239666 | - | Zm00001d035396 |
| Zm00001d001318 | 6 | 24250036 | 24250621 | + | Zm00001d001318 |
| Zm00001d035429 | 6 | 25236764 | 25238668 | + | Retrovirus-related Pol polyprotein LINE-1 |
| Zm00001d035240 | 6 | 16276743 | 16279273 | + | Zm00001d035240 |
| Zm00001d035236 | 6 | 16177948 | 16179147 | - | Plant calmodulin-binding protein-related |
| Zm00001d035235 | 6 | 16089925 | 16093478 | + | Protein cornichon homolog 1 |
| Zm00001d035233 | 6 | 15856602 | 15857420 | - | Zm00001d035233 |
| Zm00001d035234 | 6 | 15908223 | 15911138 | + | protein_coding |
| Zm00001d035232 | 6 | 15811792 | 15812517 | + | Zm00001d035232 |
| Zm00001d035227 | 6 | 15529000 | 15529392 | + | Zm00001d035227 |
| Zm00001d035228 | 6 | 15529973 | 15536315 | - | Methyl-CpG-binding domain protein 4-like protein |
| Zm00001d035229 | 6 | 15530206 | 15536423 | - | Methyl-CpG-binding domain protein 4-like protein |
| Zm00001d035224 | 6 | 14927596 | 14930694 | - | protein_coding |
| Zm00001d001304 | 6 | 14910584 | 14910813 | - | Zm00001d001304 |
| Zm00001d035323 | 6 | 21265695 | 21270357 | - | protein_coding |
| Zm00001d008563 | 8 | 13125997 | 13131966 | + | Zm00001d008563 |
| Zm00001d008564 | 8 | 13133398 | 13137232 | + | Surfeit locus protein 5 |

| | | | | | |
|---|---|---|---|---|---|
| Zm00001d008565 | 8 | 13138808 | 13143070 | + | Zm00001d008565 |
| Zm00001d025908 | 10 | 1.33E+08 | 1.33E+08 | - | OSJNBa0074L08.7 protein |
| Zm00001d025910 | 10 | 1.34E+08 | 1.34E+08 | - | protein_coding |
| Zm00001d025914 | 10 | 1.34E+08 | 1.34E+08 | + | Zm00001d025914 |
| Zm00001d025915 | 10 | 1.34E+08 | 1.34E+08 | + | Syntaxin-81 |
| Zm00001d025916 | 10 | 1.34E+08 | 1.34E+08 | - | E3 ubiquitin-protein ligase RMA1 |
| Zm00001d023177 | 10 | 1.34E+08 | 1.34E+08 | - | Zm00001d023177 |
| Zm00001d025922 | 10 | 1.34E+08 | 1.34E+08 | - | (+)-neomenthol dehydrogenase |

*Table S3: Transcripts (genes) identified using TWAS in well watered (WW) and water deficit (WD) conditions using fixed effect (FE) and random effect (RE) procedures of meta-analyses. "gene_id" represents the significant transcript. "Chr" represents chromosome number. "start" and "end" represent coordinates. "strand" represents if the gene is located on the positive or the negative strand. "TWAS_WW_FE", "TWAS_WW_RE", "TWAS_WD_FE", and "TWAS_WD_RE" represent p-values. NAs among p-values indicate that the transcript was not significant in the corresponding TWAS.*

| gene_id | Chr | start | end | strand | description | TWAS_WW_FE | TWAS_WW_RE | TWAS_WD_FE | TWAS_WD_RE |
|---|---|---|---|---|---|---|---|---|---|
| Zm00001d029973 | 1 | 97097237 | 97100691 | + | Nucleic acid binding protein | NA | 7.28E-05 | NA | NA |
| Zm00001d032131 | 1 | 2.14E+08 | 2.14E+08 | + | Scar-like domain-containing protein WAVE 5 | 2.77E-05 | NA | NA | NA |
| Zm00001d034045 | 1 | 2.82E+08 | 2.82E+08 | - | protein_coding | 4.43E-08 | 1.42E-07 | 2.24E-07 | 1.45E-05 |
| Zm00001d034047 | 1 | 2.82E+08 | 2.82E+08 | - | protein_coding | NA | 0.00010993 | NA | NA |
| Zm00001d024464 | 10 | 71966048 | 71969342 | + | Haloacid dehalogenase-like hydrolase (HAD) superfamily protein | 0.000121 | NA | NA | NA |
| Zm00001d026190 | 10 | 1.41E+08 | 1.41E+08 | - | DeSI-like protein | NA | 0.000153527 | NA | NA |
| Zm00001d002256 | 2 | 9055230 | 9066450 | + | Starch synthase 3 chloroplastic/amyloplastic | 0.000252 | 0.000108502 | NA | NA |
| Zm00001d003157 | 2 | 33981493 | 33984783 | - | protein_coding | NA | NA | 0.000297 | NA |
| Zm00001d006692 | 2 | 2.14E+08 | 2.14E+08 | - | Zm00001d006692 | 0.000206 | NA | NA | NA |
| Zm00001d007072 | 2 | 2.21E+08 | 2.21E+08 | + | Uncharacterised protein family (UPF0114) | NA | NA | 0.000182 | NA |
| Zm00001d007949 | 2 | 2.44E+08 | 2.44E+08 | + | protein_coding | NA | NA | NA | 2.14E-05 |
| Zm00001d040693 | 3 | 58561843 | 58565202 | + | Tetratricopeptide repeat (TPR)-like superfamily protein | NA | 0.000303056 | NA | NA |

| | | | | | | | | | |
|---|---|---|---|---|---|---|---|---|---|
| Zm00001d041669 | 3 | 1.33E+08 | 1.33E+08 | - | Zm00001d041669 | 0.000313 | 0.000247116 | NA | NA |
| Zm00001d041882 | 3 | 1.42E+08 | 1.42E+08 | + | protein_coding | NA | NA | 4.61E-05 | NA |
| Zm00001d043174 | 3 | 1.91E+08 | 1.91E+08 | + | protein_coding | NA | 9.91E-05 | 0.00022 | 0.000115 |
| Zm00001d043484 | 3 | 2.02E+08 | 2.02E+08 | - | Zm00001d043484 | NA | 7.33E-05 | NA | NA |
| Zm00001d044164 | 3 | 2.21E+08 | 2.21E+08 | - | Beta-adaptin-like protein A | NA | NA | 0.000287 | NA |
| Zm00001d048722 | 4 | 4187982 | 4189310 | + | Protein SHOOT GRAVITROPISM 5 | NA | 3.10E-05 | NA | NA |
| Zm00001d050918 | 4 | 1.31E+08 | 1.31E+08 | - | Zm00001d050918 | 0.000256 | NA | NA | NA |
| Zm00001d052444 | 4 | 1.9E+08 | 1.9E+08 | + | Hydroxymethylglutaryl-CoA lyase mitochondrial | NA | 0.000298524 | NA | NA |
| Zm00001d013259 | 5 | 7240613 | 7258833 | + | protein_coding | NA | NA | 0.000257 | NA |
| Zm00001d013669 | 5 | 16901324 | 16905867 | - | Chaperone protein dnaJ 3 | 0.000353 | NA | NA | NA |
| Zm00001d016530 | 5 | 1.67E+08 | 1.67E+08 | + | Probable magnesium transporter NIPA2 | 3.18E-05 | NA | NA | NA |
| Zm00001d035157 | 6 | 7246875 | 7250944 | + | Lysine histidine transporter 2 | 0.000362 | NA | NA | NA |
| Zm00001d035163 | 6 | 7409171 | 7417031 | + | Lysine histidine transporter 2 | NA | NA | 0.000178 | NA |
| Zm00001d001304 | 6 | 14910584 | 14910813 | - | Zm00001d001304 | 0.000296 | NA | NA | NA |
| Zm00001d035346 | 6 | 22716458 | 22744646 | - | Probable sucrose-phosphatase 2 | 1.02E-06 | 6.32E-13 | 9.76E-08 | 4.44E-12 |
| Zm00001d035367 | 6 | 23489630 | 23490985 | + | Zm00001d035367 | 7.67E-06 | 1.05E-11 | 6.55E-06 | 2.45E-10 |
| Zm00001d035377 | 6 | 23680168 | 23690640 | + | IQ domain-containing protein IQM3 | 2.02E-05 | 3.60E-10 | 3.57E-05 | 9.83E-08 |
| Zm00001d035382 | 6 | 23840275 | 23852509 | - | protein_coding | 0.000241 | 2.40E-06 | NA | 7.68E-07 |
| Zm00001d035390 | 6 | 24034207 | 24035363 | - | protein_coding | 2.53E-06 | 1.09E-19 | 5.49E-07 | 1.16E-20 |
| Zm00001d035400 | 6 | 24389981 | 24393065 | - | Zm00001d035400 | 0.000107 | 7.41E-06 | 2.59E-06 | 1.43E-07 |
| Zm00001d035401 | 6 | 24413321 | 24417330 | - | Putative DUF26-domain receptor-like protein kinase family protein | 2.23E-05 | 1.55E-05 | NA | 2.91E-05 |
| Zm00001d035409 | 6 | 24606673 | 24607624 | - | Abscisic stress-ripening protein 2 | 5.44E-07 | 3.26E-14 | 1.98E-05 | 1.14E-11 |
| Zm00001d035413 | 6 | 24624255 | 24626622 | + | Zm00001d035413 | 1.05E-05 | 4.95E-10 | NA | NA |
| Zm00001d035420 | 6 | 24798510 | 24811775 | + | Zm00001d035420 | 4.09E-06 | 4.63E-09 | 1.89E-05 | 7.83E-08 |

| | | | | | | | | | |
|---|---|---|---|---|---|---|---|---|---|
| Zm00001d035428 | 6 | 25148734 | 25158121 | - | Zinc finger (C3HC4-type RING finger) family protein | NA | 3.69E-05 | NA | NA |
| Zm00001d036773 | 6 | 1E+08 | 1E+08 | - | Triacylglycerol lipase 2 | NA | NA | NA | 0.000161 |
| Zm00001d022254 | 7 | 1.74E+08 | 1.74E+08 | - | Haloacid dehalogenase-like hydrolase (HAD) superfamily protein | NA | NA | 0.000179 | NA |
| Zm00001d008245 | 8 | 2490496 | 2493628 | + | protein_coding | 4.78E-06 | 5.84E-05 | 0.00022 | NA |
| Zm00001d008939 | 8 | 26134915 | 26143970 | + | protein_coding | NA | NA | NA | 0.000205 |
| Zm00001d009803 | 8 | 82156141 | 82159658 | - | NADH-ubiquinone oxidoreductase 10.5 kDa subunit%253B NADH-ubiquinone oxidoreductase subunit | 1.73E-05 | 2.02E-05 | NA | NA |
| Zm00001d010752 | 8 | 1.27E+08 | 1.27E+08 | - | protein_coding | 1.11E-05 | NA | NA | NA |
| Zm00001d011882 | 8 | 1.64E+08 | 1.64E+08 | + | protein_coding | 7.69E-05 | 5.88E-05 | NA | NA |
| Zm00001d011904 | 8 | 1.64E+08 | 1.64E+08 | + | Vesicle-associated protein 2-2 | 0.00042 | NA | NA | NA |
| Zm00001d012144 | 8 | 1.69E+08 | 1.69E+08 | - | 4-coumarate--CoA ligase-like 9 | NA | NA | 8.46E-05 | 0.000139 |
| Zm00001d045205 | 9 | 16041474 | 16043191 | + | protein_coding | 0.000157 | 0.00019538 | 0.000162 | NA |

*Table S4: Significant SNPs identified using eQTL analyses on WW biomarkers (significant transcripts identified in TWAS). "SNP" represents the name of the SNP. "pval" represents eQTL analysis p-value, "snp_chr" represents the chromosome on which the SNP is present. "eQTL" represents if the SNP is a part of Cis or Trans eQTL. "gene" represents the transcript for which the SNP was identified. "gene_chr" represents the chromosome on which the transcript is located. The last columns are the positions of the SNP and gene in Mb, and the distance between the SNP and the gene when they are on the same chromosome.*

| SNP | pval | gene | eQTL | eqtl_description | snp_chr | gene_chr | snp_pos_Mb | gene_start_Mb | gene_end_Mb | dist_start_Mb | dist_end_Mb |
|---|---|---|---|---|---|---|---|---|---|---|---|
| AX-90982390 | 1,04E-22 | Zm00001d001304 | Cis | Downstream | 6 | 6 | 15.227158 | 14.910584 | 14.910813 | 0.316574 | 0.316345 |
| AX-90922003 | 3,86E-20 | Zm00001d006692 | Trans | Different Chr | 5 | 2 | 5.510533 | 214.487406 | 214.491116 | NA | NA |
| AX-90553036 | 3,88E-25 | Zm00001d008245 | Cis | Inside Gene | 8 | 8 | 2.493622 | 2.490496 | 2.493628 | 0.003126 | -6,00E-06 |
| AX-91089088 | 2,46E-14 | Zm00001d009803 | Cis | Downstream | 8 | 8 | 82.394389 | 82.156141 | 82.159658 | 0.238248 | 0.234731 |
| S9_7203689 | 2,31E-09 | Zm00001d010752 | Trans | Different Chr | 9 | 8 | 6.789554 | 126.880531 | 126.882389 | NA | NA |
| AX-91385006 | 3,68E-08 | Zm00001d010752 | Trans | Different Chr | 5 | 8 | 0.984724 | 126.880531 | 126.882389 | NA | NA |
| AX-91110629 | 1,47E-19 | Zm00001d011882 | Cis | Inside Gene | 8 | 8 | 163.877227 | 163.875069 | 163.87871 | 0.002158 | -0.001483 |
| S1_207527452 | 7,35E-15 | Zm00001d011882 | Trans | Different Chr | 1 | 8 | 210.344428 | 163.875069 | 163.87871 | NA | NA |
| AX-90963076 | 4,28E-12 | Zm00001d016530 | Cis | Inside Gene | 5 | 5 | 166.562644 | 166.555478 | 166.56355 | 0.007166 | -0.000906 |

| | | | | | | | | | | | |
|---|---|---|---|---|---|---|---|---|---|---|---|
| AX-91176081 | 2,73E-10 | Zm00001d024464 | Cis | Upstream | 10 | 10 | 71.8652 | 71.966048 | 71.969342 | -0.100848 | -0.104142 |
| AX-91193782 | 5,42E-13 | Zm00001d026190 | Cis | Downstream | 10 | 10 | 140.56521 | 140.561973 | 140.56517 | 0.003237 | 4,00E-05 |
| S10_139942871 | 4,36E-09 | Zm00001d026190 | Cis | Downstream | 10 | 10 | 140.6452 | 140.561973 | 140.56517 | 0.083227 | 0.08003 |
| S1_93970703 | 7,81E-22 | Zm00001d029973 | Cis | Upstream | 1 | 1 | 96.146752 | 97.097237 | 97.100691 | -0.950485 | -0.953939 |
| AX-90615167 | 1,05E-15 | Zm00001d029973 | Cis | Inside Gene | 1 | 1 | 97.100327 | 97.097237 | 97.100691 | 0.00309 | -0.000364 |
| S5_3426248 | 4,69E-08 | Zm00001d029973 | Trans | Different Chr | 5 | 1 | 3.490341 | 97.097237 | 97.100691 | NA | NA |
| AX-90704705 | 1,31E-62 | Zm00001d032131 | Cis | Upstream | 1 | 1 | 213.573451 | 213.971572 | 213.974101 | -0.398121 | -0.40065 |
| S6_5644206 | 6,35E-18 | Zm00001d035157 | Trans | NA | 6 | 6 | 6.130471 | 7.246875 | 7.250944 | -1.116404 | -1.120473 |
| AX-90982330 | 5,16E-131 | Zm00001d035346 | Trans | NA | 6 | 6 | 15.535341 | 22.716458 | 22.744646 | -7.181117 | -7.209305 |
| S8_170864043 | 1,47E-12 | Zm00001d035346 | Trans | Different Chr | 8 | 6 | 176.105298 | 22.716458 | 22.744646 | NA | NA |
| S1_200629103 | 4,43E-08 | Zm00001d035346 | Trans | Different Chr | 1 | 6 | 203.1279 | 22.716458 | 22.744646 | NA | NA |
| AX-90982330 | 9,07E-121 | Zm00001d035367 | Trans | NA | 6 | 6 | 15.535341 | 23.48963 | 23.490985 | -7.954289 | -7.955644 |
| AX-91683421 | 4,62E-10 | Zm00001d035367 | Trans | NA | 6 | 6 | 8.541559 | 23.48963 | 23.490985 | -14.948071 | -14.949426 |
| AX-90982330 | 2,31E-60 | Zm00001d035377 | Trans | NA | 6 | 6 | 15.535341 | 23.680168 | 23.69064 | -8.144827 | -8.155299 |
| S6_18528943 | 1,59E-13 | Zm00001d035377 | Trans | NA | 6 | 6 | 16.285728 | 23.680168 | 23.69064 | -7.39444 | -7.404912 |
| AX-91352977 | 7,98E-09 | Zm00001d035377 | Trans | NA | 6 | 6 | 10.333075 | 23.680168 | 23.69064 | -13.347093 | -13.357565 |
| AX-90982330 | 5,79E-45 | Zm00001d035382 | Trans | NA | 6 | 6 | 15.535341 | 23.840275 | 23.852509 | -8.304934 | -8.317168 |
| AX-90981217 | 1,46E-42 | Zm00001d035390 | Cis | Downstream | 6 | 6 | 24.449216 | 24.034207 | 24.035363 | 0.415009 | 0.413853 |
| PZE-106021419 | 4,68E-11 | Zm00001d035390 | Trans | NA | 6 | 6 | 15.812017 | 24.034207 | 24.035363 | -8.22219 | -8.223346 |
| S6_32887134 | 7,23E-16 | Zm00001d035390 | Trans | NA | 6 | 6 | 34.422825 | 24.034207 | 24.035363 | 10.388618 | 10.387462 |
| AX-90787663 | 8,07E-10 | Zm00001d035390 | Trans | Different Chr | 2 | 6 | 222.134193 | 24.034207 | 24.035363 | NA | NA |
| S6_18527553 | 1,32E-10 | Zm00001d035390 | Trans | NA | 6 | 6 | 16.287118 | 24.034207 | 24.035363 | -7.747089 | -7.748245 |
| S6_25289719 | 4,43E-08 | Zm00001d035390 | Trans | NA | 6 | 6 | 26.665542 | 24.034207 | 24.035363 | 2.631335 | 2.630179 |
| AX-90982390 | 2,28E-46 | Zm00001d035400 | Trans | NA | 6 | 6 | 15.227158 | 24.389981 | 24.393065 | -9.162823 | -9.165907 |
| AX-90982390 | 3,24E-34 | Zm00001d035401 | Trans | NA | 6 | 6 | 15.227158 | 24.413321 | 24.41733 | -9.186163 | -9.190172 |
| S6_14852888 | 2,91E-15 | Zm00001d035401 | Cis | Downstream | 6 | 6 | 24.705761 | 24.413321 | 24.41733 | 0.29244 | 0.288431 |
| AX-90981217 | 1,54E-21 | Zm00001d035409 | Cis | Upstream | 6 | 6 | 24.449216 | 24.606673 | 24.607624 | -0.157457 | -0.158408 |
| AX-90980434 | 1,18E-10 | Zm00001d035409 | Trans | NA | 6 | 6 | 9.904391 | 24.606673 | 24.607624 | -14.702282 | -14.703233 |
| AX-90982107 | 6,14E-42 | Zm00001d035409 | Trans | NA | 6 | 6 | 16.574536 | 24.606673 | 24.607624 | -8.032137 | -8.033088 |
| AX-90981486 | 2,61E-29 | Zm00001d035409 | Cis | Downstream | 6 | 6 | 25.279456 | 24.606673 | 24.607624 | 0.672783 | 0.671832 |
| AX-90598121 | 1,29E-11 | Zm00001d035409 | Trans | Different Chr | 8 | 6 | 174.965618 | 24.606673 | 24.607624 | NA | NA |
| S5_139961666 | 6,57E-10 | Zm00001d035409 | Trans | Different Chr | 5 | 6 | 143.333467 | 24.606673 | 24.607624 | NA | NA |

| | | | | | | | | | | | |
|---|---|---|---|---|---|---|---|---|---|---|---|
| S3_34388439 | 1,05E-08 | Zm00001d035409 | Trans | Different Chr | 3 | 6 | 34.080299 | 24.606673 | 24.607624 | NA | NA |
| AX-90981217 | 2,96E-33 | Zm00001d035413 | Cis | Upstream | 6 | 6 | 24.449216 | 24.624255 | 24.626622 | -0.175039 | -0.177406 |
| AX-91684390 | 8,00E-41 | Zm00001d035413 | Trans | NA | 6 | 6 | 18.16695 | 24.624255 | 24.626622 | -6.457305 | -6.459672 |
| S6_18528881 | 6,90E-46 | Zm00001d035413 | Trans | NA | 6 | 6 | 16.28579 | 24.624255 | 24.626622 | -8.338465 | -8.340832 |
| S6_21888221 | 3,27E-15 | Zm00001d035413 | Trans | NA | 6 | 6 | 20.190487 | 24.624255 | 24.626622 | -4.433768 | -4.436135 |
| S6_4614215 | 5,61E-15 | Zm00001d035413 | Trans | NA | 6 | 6 | 4.659631 | 24.624255 | 24.626622 | -19.964624 | -19.966991 |
| AX-91416291 | 3,84E-08 | Zm00001d035413 | Trans | Different Chr | 7 | 6 | 116.463658 | 24.624255 | 24.626622 | NA | NA |
| AX-90982330 | 9,01E-102 | Zm00001d035420 | Trans | NA | 6 | 6 | 15.535341 | 24.79851 | 24.811775 | -9.263169 | -9.276434 |
| AX-90982024 | 1,30E-14 | Zm00001d035428 | Trans | NA | 6 | 6 | 16.820991 | 25.148734 | 25.158121 | -8.327743 | -8.33713 |
| AX-91847705 | 2,37E-42 | Zm00001d040693 | Cis | Downstream | 3 | 3 | 58.892325 | 58.561843 | 58.565202 | 0.330482 | 0.327123 |
| AX-91413128 | 1,88E-08 | Zm00001d040693 | Trans | Different Chr | 8 | 3 | 25.304493 | 58.561843 | 58.565202 | NA | NA |
| PZE-106059327 | 2,82E-19 | Zm00001d041669 | Trans | Different Chr | 6 | 3 | 112.863316 | 132.743387 | 132.744631 | NA | NA |
| AX-91646937 | 3,73E-26 | Zm00001d041669 | Trans | Different Chr | 5 | 3 | 27.979136 | 132.743387 | 132.744631 | NA | NA |
| S3_159911470 | 3,67E-21 | Zm00001d041669 | Trans | NA | 3 | 3 | 161.506509 | 132.743387 | 132.744631 | 28.763122 | 28.761878 |
| S5_213916149 | 1,17E-20 | Zm00001d041669 | Trans | Different Chr | 5 | 3 | 219.785384 | 132.743387 | 132.744631 | NA | NA |
| AX-91083340 | 1,51E-24 | Zm00001d041669 | Trans | Different Chr | 8 | 3 | 59.484108 | 132.743387 | 132.744631 | NA | NA |
| AX-90588310 | 7,27E-16 | Zm00001d041669 | Trans | Different Chr | 6 | 3 | 117.125033 | 132.743387 | 132.744631 | NA | NA |
| S8_28852380 | 4,51E-11 | Zm00001d041669 | Trans | Different Chr | 8 | 3 | 29.78929 | 132.743387 | 132.744631 | NA | NA |
| S9_6997871 | 4,49E-08 | Zm00001d041669 | Trans | Different Chr | 9 | 3 | 6.586426 | 132.743387 | 132.744631 | NA | NA |
| S3_187711137 | 1,65E-31 | Zm00001d043174 | Cis | Inside Gene | 3 | 3 | 190.626036 | 190.625724 | 190.628445 | 0.000312 | -0.002409 |
| S3_187656231 | 1,20E-19 | Zm00001d043174 | Cis | Upstream | 3 | 3 | 190.577309 | 190.625724 | 190.628445 | -0.048415 | -0.051136 |
| AX-90845736 | 1,42E-97 | Zm00001d043484 | Cis | Upstream | 3 | 3 | 201.604343 | 201.62493 | 201.630525 | -0.020587 | -0.026182 |
| S7_143036001 | 3,55E-10 | Zm00001d043484 | Trans | Different Chr | 7 | 3 | 147.986326 | 201.62493 | 201.630525 | NA | NA |
| AX-91436364 | 3,21E-25 | Zm00001d045205 | Cis | Inside Gene | 9 | 9 | 16.043021 | 16.041474 | 16.043191 | 0.001547 | -0.00017 |
| S4_3483048 | 2,44E-23 | Zm00001d048722 | Cis | Upstream | 4 | 4 | 4.187469 | 4.187982 | 4.18931 | -0.000513 | -0.001841 |
| AX-90579324 | 5,20E-08 | Zm00001d048722 | Cis | Downstream | 4 | 4 | 4.48395 | 4.187982 | 4.18931 | 0.295968 | 0.29464 |
| AX-91621026 | 3,59E-22 | Zm00001d050918 | Cis | Upstream | 4 | 4 | 130.987651 | 130.988293 | 130.988613 | -0.000642 | -0.000962 |
| AX-91422067 | 7,53E-17 | Zm00001d052444 | Cis | Downstream | 4 | 4 | 190.148986 | 190.133615 | 190.136497 | 0.015371 | 0.012489 |

*Table S5 : Significant SNPs identified using eQTL analyses on WD biomarkers (significant transcripts identified in TWAS). "SNP" represents the name of the SNP. "pval" represents eQTL analysis p-value, "snp_chr" represents the chromosome on which the SNP is present. "eQTL" represents if the SNP is a part of Cis or Trans eQTL. "gene" represents the transcript for which the SNP was identified. "gene_chr" represents the chromosome on which the transcript is located. The last columns are the positions of the SNP and gene in Mb, and the distance between the SNP and the gene when they are on the same chromosome.*

| SNP | pval | gene | eQTL | eqtl_description | snp_chr | gene_chr | snp_pos_Mb | gene_start_Mb | gene_end_Mb | dist_start_Mb | dist_end_Mb |
|---|---|---|---|---|---|---|---|---|---|---|---|
| AX-90739517 | 1,62E-09 | Zm00001d003157 | Trans | NA | 2 | 2 | 35.310259 | 33.981493 | 33.984783 | 1.328766 | 1.325476 |
| AX-90629050 | 4,42E-36 | Zm00001d007072 | Cis | Inside Gene | 2 | 2 | 221.454914 | 221.453612 | 221.455343 | 0.001302 | -0.000429 |
| S2_214661112 | 6,77E-13 | Zm00001d007072 | Cis | Upstream | 2 | 2 | 221.449244 | 221.453612 | 221.455343 | -0.004368 | -0.006099 |
| S2_214661266 | 1,67E-10 | Zm00001d007072 | Cis | Upstream | 2 | 2 | 221.449398 | 221.453612 | 221.455343 | -0.004214 | -0.005945 |
| AX-90553036 | 7,68E-30 | Zm00001d008245 | Cis | Inside Gene | 8 | 8 | 2.493622 | 2.490496 | 2.493628 | 0.003126 | -6,00E-06 |
| PZE-103050856 | 2,27E-08 | Zm00001d008245 | Trans | Different Chr | 3 | 8 | 55.960426 | 2.490496 | 2.493628 | NA | NA |
| AX-91390970 | 5,00E-21 | Zm00001d012144 | Cis | Downstream | 8 | 8 | 168.871721 | 168.855687 | 168.867132 | 0.016034 | 0.004589 |
| AX-90974802 | 8,06E-12 | Zm00001d013259 | Trans | NA | 5 | 5 | 211.812194 | 7.240613 | 7.258833 | 204.571581 | 204.553361 |
| PZE-103087869 | 5,32E-09 | Zm00001d013259 | Trans | Different Chr | 3 | 5 | 146.923108 | 7.240613 | 7.258833 | NA | NA |
| S1_148515009 | 2,46E-10 | Zm00001d013259 | Trans | Different Chr | 1 | 5 | 150.364875 | 7.240613 | 7.258833 | NA | NA |
| S1_191581451 | 5,01E-09 | Zm00001d013259 | Trans | Different Chr | 1 | 5 | 193.835824 | 7.240613 | 7.258833 | NA | NA |
| AX-91743591 | 2,72E-12 | Zm00001d022254 | Cis | Downstream | 7 | 7 | 173.716594 | 173.713829 | 173.715713 | 0.002765 | 0.000881 |
| S7_168218260 | 4,84E-10 | Zm00001d022254 | Cis | Downstream | 7 | 7 | 173.754387 | 173.713829 | 173.715713 | 0.040558 | 0.038674 |
| AX-90979946 | 1,58E-19 | Zm00001d035163 | Cis | Inside Gene | 6 | 6 | 7.414245 | 7.409171 | 7.417031 | 0.005074 | -0.002786 |
| AX-90630357 | 9,21E-122 | Zm00001d035346 | Trans | NA | 6 | 6 | 15.530801 | 22.716458 | 22.744646 | -7.185657 | -7.213845 |
| AX-91545917 | 2,50E-08 | Zm00001d035346 | Trans | Different Chr | 2 | 6 | 200.423165 | 22.716458 | 22.744646 | NA | NA |
| AX-90630357 | 3,51E-124 | Zm00001d035367 | Trans | NA | 6 | 6 | 15.530801 | 23.48963 | 23.490985 | -7.958829 | -7.960184 |
| AX-91683421 | 4,48E-09 | Zm00001d035367 | Trans | NA | 6 | 6 | 8.541559 | 23.48963 | 23.490985 | -14.948071 | -14.949426 |
| PZE-106021419 | 1,49E-84 | Zm00001d035377 | Trans | NA | 6 | 6 | 15.812017 | 23.680168 | 23.69064 | -7.868151 | -7.878623 |
| AX-91352977 | 5,53E-16 | Zm00001d035377 | Trans | NA | 6 | 6 | 10.333075 | 23.680168 | 23.69064 | -13.347093 | -13.357565 |

| | | | | | | | | | | | |
|---|---|---|---|---|---|---|---|---|---|---|---|
| S6_18528931 | 8,55E-12 | Zm00001d035377 | Trans | NA | 6 | 6 | 16.28574 | 23.680168 | 23.69064 | -7.394428 | -7.4049 |
| S1_2753482 | 1,23E-08 | Zm00001d035377 | Trans | Different Chr | 1 | 6 | 2.895415 | 23.680168 | 23.69064 | NA | NA |
| AX-90982316 | 9,14E-48 | Zm00001d035382 | Trans | NA | 6 | 6 | 15.531892 | 23.840275 | 23.852509 | -8.308383 | -8.320617 |
| AX-90548528 | 1,05E-48 | Zm00001d035390 | Cis | Downstream | 6 | 6 | 24.261441 | 24.034207 | 24.035363 | 0.227234 | 0.226078 |
| AX-90982269 | 2,94E-20 | Zm00001d035390 | Trans | NA | 6 | 6 | 15.810754 | 24.034207 | 24.035363 | -8.223453 | -8.224609 |
| S6_32967902 | 1,10E-12 | Zm00001d035390 | Trans | NA | 6 | 6 | 34.510046 | 24.034207 | 24.035363 | 10.475839 | 10.474683 |
| S10_145097698 | 1,63E-09 | Zm00001d035390 | Trans | Different Chr | 10 | 6 | 145.798453 | 24.034207 | 24.035363 | NA | NA |
| S1_260932507 | 1,18E-08 | Zm00001d035390 | Trans | Different Chr | 1 | 6 | 265.765418 | 24.034207 | 24.035363 | NA | NA |
| AX-90622419 | 7,72E-64 | Zm00001d035400 | Trans | NA | 6 | 6 | 15.534324 | 24.389981 | 24.393065 | -8.855657 | -8.858741 |
| S5_76381740 | 6,69E-10 | Zm00001d035400 | Trans | Different Chr | 5 | 6 | 78.676259 | 24.389981 | 24.393065 | NA | NA |
| AX-91071026 | 1,91E-09 | Zm00001d035400 | Trans | Different Chr | 8 | 6 | 8.465247 | 24.389981 | 24.393065 | NA | NA |
| AX-91652195 | 3,12E-09 | Zm00001d035400 | Trans | Different Chr | 5 | 6 | 57.157036 | 24.389981 | 24.393065 | NA | NA |
| AX-91504865 | 1,67E-10 | Zm00001d035400 | Trans | Different Chr | 1 | 6 | 283.458778 | 24.389981 | 24.393065 | NA | NA |
| S10_127360125 | 4,36E-09 | Zm00001d035400 | Trans | Different Chr | 10 | 6 | 128.390471 | 24.389981 | 24.393065 | NA | NA |
| S1_174285814 | 5,08E-08 | Zm00001d035400 | Trans | Different Chr | 11 | 6 | 0.034159 | 24.389981 | 24.393065 | NA | NA |
| PZE-106021419 | 2,18E-67 | Zm00001d035401 | Trans | NA | 6 | 6 | 15.812017 | 24.413321 | 24.41733 | -8.601304 | -8.605313 |
| AX-90983381 | 1,74E-19 | Zm00001d035401 | Trans | NA | 6 | 6 | 25.569241 | 24.413321 | 24.41733 | 1.15592 | 1.151911 |
| AX-90981217 | 1,18E-40 | Zm00001d035409 | Cis | Upstream | 6 | 6 | 24.449216 | 24.606673 | 24.607624 | -0.157457 | -0.158408 |
| PZE-106021183 | 1,17E-80 | Zm00001d035409 | Trans | NA | 6 | 6 | 16.572658 | 24.606673 | 24.607624 | -8.034015 | -8.034966 |
| AX-90981486 | 7,24E-60 | Zm00001d035409 | Cis | Downstream | 6 | 6 | 25.279456 | 24.606673 | 24.607624 | 0.672783 | 0.671832 |
| S4_228666403 | 4,06E-28 | Zm00001d035409 | Trans | Different Chr | 4 | 6 | 233.64169 | 24.606673 | 24.607624 | NA | NA |
| S1_195558459 | 6,65E-18 | Zm00001d035409 | Trans | Different Chr | 1 | 6 | 197.918511 | 24.606673 | 24.607624 | NA | NA |
| AX-90982407 | 1,62E-11 | Zm00001d035409 | Trans | NA | 6 | 6 | 15.114737 | 24.606673 | 24.607624 | -9.491936 | -9.492887 |

| | | | | | | | | | | | |
|---|---|---|---|---|---|---|---|---|---|---|---|
| AX-90998799 | 2,02E-15 | Zm00001d035409 | Trans | NA | 6 | 6 | 89.081991 | 24.606673 | 24.607624 | 64.475318 | 64.474367 |
| S2_15994974 | 6,95E-12 | Zm00001d035409 | Trans | Different Chr | 2 | 6 | 16.417097 | 24.606673 | 24.607624 | NA | NA |
| S3_221694783 | 9,03E-09 | Zm00001d035409 | Trans | Different Chr | 3 | 6 | 225.46426 | 24.606673 | 24.607624 | NA | NA |
| AX-90630357 | 6,54E-114 | Zm00001d035420 | Trans | NA | 6 | 6 | 15.530801 | 24.79851 | 24.811775 | -9.267709 | -9.280974 |
| S6_102756416 | 2,33E-60 | Zm00001d036773 | Trans | NA | 6 | 6 | 106.002954 | 100.236033 | 100.238454 | 5.766921 | 5.7645 |
| AX-91452330 | 1,24E-11 | Zm00001d036773 | Trans | Different Chr | 4 | 6 | 234.971287 | 100.236033 | 100.238454 | NA | NA |
| S7_131034746 | 3,07E-09 | Zm00001d036773 | Trans | Different Chr | 7 | 6 | 135.065097 | 100.236033 | 100.238454 | NA | NA |
| AX-91580208 | 6,68E-13 | Zm00001d041882 | Cis | Inside Gene | 3 | 3 | 142.231454 | 142.230896 | 142.233815 | 0.000558 | -0.002361 |
| S3_187898809 | 1,12E-33 | Zm00001d043174 | Cis | Downstream | 3 | 3 | 190.786283 | 190.625724 | 190.628445 | 0.160559 | 0.157838 |
| S3_187656231 | 1,58E-28 | Zm00001d043174 | Cis | Upstream | 3 | 3 | 190.577309 | 190.625724 | 190.628445 | -0.048415 | -0.051136 |
| AX-91166155 | 3,79E-09 | Zm00001d043174 | Trans | Different Chr | 10 | 3 | 36.875691 | 190.625724 | 190.628445 | NA | NA |
| S3_195921881 | 3,90E-09 | Zm00001d043174 | Trans | NA | 3 | 3 | 198.902598 | 190.625724 | 190.628445 | 8.276874 | 8.274153 |
| S3_217237862 | 3,59E-21 | Zm00001d044164 | Cis | Inside Gene | 3 | 3 | 220.849966 | 220.843094 | 220.855486 | 0.006872 | -0.00552 |
| AX-91436364 | 3,15E-25 | Zm00001d045205 | Cis | Inside Gene | 9 | 9 | 16.043021 | 16.041474 | 16.043191 | 0.001547 | -0.00017 |
| S9_16997589 | 3,14E-08 | Zm00001d045205 | Cis | Downstream | 9 | 9 | 16.751173 | 16.041474 | 16.043191 | 0.709699 | 0.707982 |
